# The distinct effects of MEK and GSK3 inhibition upon the methylome and transcriptome of mouse embryonic stem cells

**DOI:** 10.1101/2021.11.18.469000

**Authors:** Julia Spindel, Christel Krueger, Felix Krueger, Evangelia K. Papachristou, Kamal Kishore, Clive S. D’Santos, Wolf Reik

## Abstract

Mouse embryonic stem cells (mESCs) were first cultured *in vitro* in serum-containing medium with leukaemia inhibitory factor, in which they exhibit heterogeneous expression of both pluripotency and some early differentiation markers. Over the last decade, however, it has become commonplace to grow mESCs with inhibitors of MEK and GSK3 signalling, which together elicit a more homogeneously ‘naive’ state of pluripotency. Whilst 2i/L-cultured mESCs have been shown to be globally hypomethylated, a comprehensive understanding of the distinct effects of these signalling inhibitors upon the DNA methylome is still lacking. Here we carried out whole genome bisulphite and RNA sequencing of mESCs grown with MEK or GSK3 inhibition alone, including different time points and concentrations of MEK inhibitor treatment. This demonstrated that MEK inhibition causes a dose-dependent impairment of maintenance methylation via loss of UHRF1 protein, as well as rapid impairment of *de novo* methylation. In contrast, GSK3 inhibition triggers impairment of *de novo* methylation alone, and consequent hypomethylation is enriched at enhancers with a 2i/L-specific chromatin signature and coincides with upregulation of nearby genes. Our study provides detailed insights into the epigenetic and transcriptional impacts of inhibiting MEK or GSK3 signalling in mouse pluripotent cells.

**Highlights:**

- MEK inhibition causes dose-dependent impairment of maintenance methylation via loss of UHRF1 protein, as well as impairment of *de novo* methylation.
- GSK3 inhibition triggers impairment of *de novo* methylation alone, which results in hypomethylation of enhancers and non-CGI promoters.
- Enhancers that are hypomethylated following GSK3 inhibition are enriched for 2i/L-specific pluripotency factor binding, TET2 and H3K4me1.
- Enhancer hypomethylation coincides with increased expression of nearby and Capture Hi-C linked genes.

## Introduction

The first successful derivation of mouse embryonic stem cells (mESCs) from explanted blastocysts required their culture atop a layer of irradiated mouse embryonic fibroblasts (MEFs) in a serum-containing medium (Evans and Kaufman, 1981; Martin, 1981). It was subsequently discovered that culture with cytokine leukaemia inhibitory factor (LIF), on an extracellular matrix such as gelatin, abolished mESC dependency on MEFs but not upon serum in the culture media (Smith et al., 1988). mESCs grown in this serum- and LIF-containing medium (S/L) are heterogeneous morphologically, transcriptomically, as well as in their protein expression of key pluripotency and some early differentiation markers (Chambers et al., 2007; Hayashi et al., 2008; Toyooka et al., 2008). Culture with inhibitors of MEK and GSK3 signalling, however, removes the need for serum in the culture medium and renders mESCs much more homogeneously reminiscent of the E4.5 epiblast (Marks et al., 2012; Wray et al., 2010; Ying et al., 2008). MEK inhibition impedes differentiation-inducing signalling from FGF4, whilst GSK3 inhibition promotes self-renewal and viability under these conditions as well as antagonising TCF3 repression of pluripotency-related genes (Kunath et al., 2007; Wray et al., 2011; Ying et al., 2008). mESCs grown in this chemically defined, serum-free, two inhibitor- and LIF-containing medium (2i/L) lack lineage bias and therefore are commonly referred to as naïve, or ground state, pluripotent stem cells.

2i/L-cultured mESCs also represent an epigenetic blank slate relative to S/L-cultured cells. They have low levels of DNA methylation throughout the genome, reduced H3K27me3 at promoters, and exhibit fewer bivalent chromatin domains overall (Ficz et al., 2013; Habibi et al., 2013; Marks et al., 2012). *De novo* methyltransferases *Dnmt3a, Dnmt3b* and *Dnmt3l* are transcriptionally downregulated in mESCs cultured in 2i/L, shown to be regulated by transcription factor PRDM14 (Ficz et al., 2013; Hackett et al., 2013; Leitch et al., 2013; Yamaji et al., 2013).

PRDM14 also mediates methylation of DNMT3A/B proteins by G9a/GLP, which in turn regulates DNMT3A/B protein stability (Sim et al., 2017). Furthermore, a key cofactor of maintenance methyltransferase DNMT1, UHRF1, is downregulated specifically at the protein level in 2i/L, and may also be less efficiently recruited to chromatin due to a simultaneous reduction of H3K9me2 (von Meyenn et al., 2016). Several H3K9 demethylases are upregulated at the protein level in 2i/L, and JMJD2C (KDM4C) is reportedly also stabilised by MEK inhibition (Sim et al., 2017; von Meyenn et al., 2016).

Whilst the cellular identity of 2i/L-cultured mESCs has been well characterised, a comprehensive understanding of the effects that MEK and GSK3 inhibitors have individually upon the epigenome and transcriptome is lacking. Investigation of this would not only improve our understanding of the establishment and maintenance of naïve pluripotency, but also of the roles of MEK and GSK3 signalling in mESCs more generally. With this aim, we carried out whole genome bisulphite and RNA sequencing, as well as whole proteome analysis, of mESCs grown with either MEK or GSK3 inhibition. Whilst MEK inhibition caused a dose-dependent impairment of maintenance methylation as well as rapid impairment of *de novo* methylation, GSK3 inhibition triggered impairment of *de novo* methylation only, providing a physiological system in which to investigate the epigenetic and transcriptomic changes associated with modulation of *de novo* methylation.

## Results

### MEK inhibition causes dose-dependent impairment of maintenance methylation

In order to assess the effects of inhibition of MEK or GSK3 signalling individually, we cultured mESCs for 8 days in either S/L, 2i/L, GSK3i/L (same basal media as 2i/L, plus LIF, but with GSK3 inhibition only) or MEKi/L (same basal media as 2i/L, plus LIF, but with MEK inhibition only), as well as for 24 days in GSK3i/L to evaluate longer-term impacts of this (Fig. S1A). We first interrogated global DNA methylation levels across these different culture conditions. As expected, cells maintained in S/L conditions retained high levels of global methylation. Whilst culture for either 8 or 24 days in GSK3i/L resulted in very marginal genome-wide demethylation, culture in MEKi/L triggered extensive hypomethylation, more pronounced even than that in 2i/L. This substantiates the previous observation that MEK inhibitor added to chemically ill-defined S/L media induces global DNA hypomethylation, and that Mek1/2, but not Gsk3a/b, double-knockout mESCs are globally hypomethylated in S/L (Choi et al., 2017). Methylation levels across genes, intergenic regions and non-CpG island promoters were consistent with genome-wide levels, whilst CpG islands remained hypomethylated across all conditions as expected (Fig. 1A, S1B & S1C). Methylation of major repeat classes also mirrored genome-wide patterns, whilst MMERVK10C, IAP and RLTR elements were resistant to demethylation in both MEKi/L and 2i/L, as previously shown for other globally demethylating systems (Berrens et al., 2017; Kobayashi et al., 2013; Seisenberger et al., 2012) (Fig. S1D).

**Figure 1:**
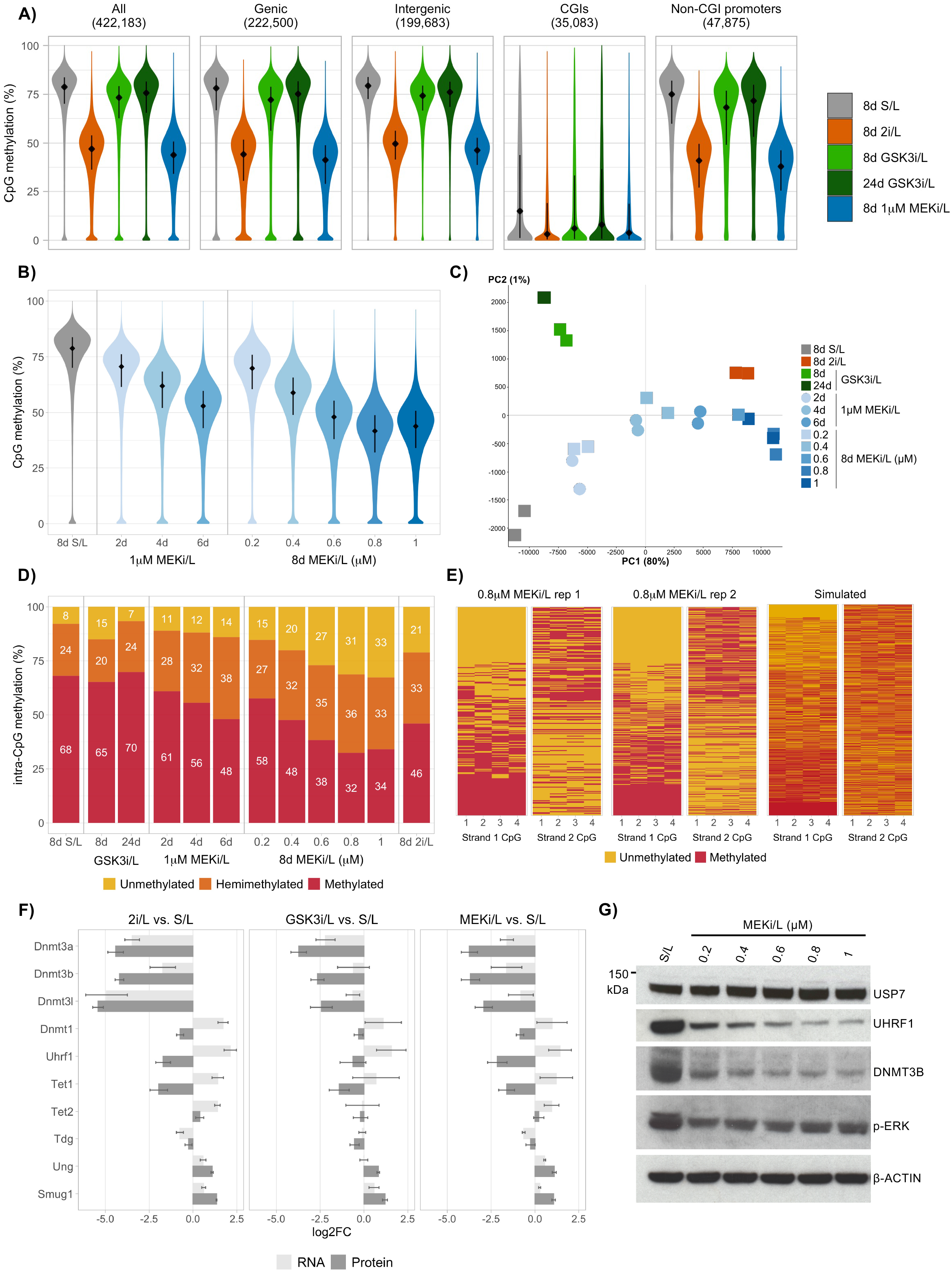
MEK inhibition causes dose-dependent impairment of maintenance methylation. **A)** Violin plots depicting genome-wide DNA methylation levels across culture conditions, as measured by WGBS. Percentage methylation was quantified over all 100 CpG windows with at least 20 observations (All), those overlapping genes (Genic), those not overlapping genes (Intergenic), those overlapping CpG islands (CGIs), and those overlapping promoters (− 2000bp to +500bp around the transcription start site) but not overlapping CGIs (Non-CGI promoters). The panels are labelled with the number of windows in each group. **B)** Violin plots depicting genome-wide DNA methylation levels across MEKi/L culture conditions, as measured by WGBS. Percentage methylation was quantified over all 100 CpG windows with at least 20 observations. **C)** Principal component analysis of individual biological replicates based on genome-wide DNA methylation levels quantified over 100 CpG windows with at least 20 observations. **D)** Percentages of unmethylated, hemimethylated and fully methylated CpG/CpG dyads across culture conditions, calculated using *in silico* strand annealing. **E)** Heatmaps of CpG methylation status for reads with ≥3 hemimethylated CpG dyads, illustrating the patterning of methylation on opposing strands. The first two heatmaps show this for biological replicates of cells grown in 0.8µM MEKi/L for 8 days, the third for simulated data for which each CpG has a 50% chance of being methylated and reads with ≥3 hemimethylated CpGs were selected. Heatmaps are ordered by the total number of methylated CpGs in Strand 1. **F)** Log2-transformed fold change of RNA and protein abundance of DNA methylation machinery in 8d 2i/L, GSK3i/L and MEKi/L culture conditions, each compared to S/L. RNA abundance was measured by RNA-seq and counts normalised per million mapped reads (RPM); protein abundance was measured by whole proteome TMT mass spectrometry. Error bars show standard deviation. **G)** Western blot showing the abundance of USP7, UHRF1, DNMT3B and phosphorylated ERK proteins in MEKi/L culture conditions with titrated concentrations of MEK inhibitor. β-ACTIN quantity is shown as a loading control.

Genome-wide demethylation progressed throughout the 8-day (8d) culture in MEKi/L, with most of the genome losing methylation gradually rather than distinct sets of regions becoming hypomethylated at particular time points. Strikingly, culture for 8 days in different concentrations of MEK inhibitor within MEKi/L demonstrated that resulting global hypomethylation was dose-dependent (Fig. 1B & S1E). Principal component analysis of DNA methylation data reveals a trajectory of global demethylation along PC1, which captures 80% of the variation across individual samples and separates S/L from 2i/L samples. 2d 1µM MEKi/L samples cluster most closely with 8d 0.2µM MEKi/L samples, 4d 1µM MEKi/L samples with 8d 0.4µM MEKi/L samples, and 6d 1µM MEKi/L samples with 8d 0.6µM MEKi/L samples. GSK3i/L samples are separated from all others along PC2 (Fig. 1C).

We performed *in silico* strand annealing to investigate the levels of hemimethylation across our samples (Xu and Corces, 2019). CpG hemimethylation levels were increased with longer culture in MEKi/L up to 6 days, or with higher concentrations of MEK inhibitor within MEKi/L up to 0.8µM, with samples cultured for 6 days in 1µM MEKi/L exhibiting 38% CpG hemimethylation (Fig. 1D). Filtering of reads with at least 3 hemimethylated CpG dyads revealed a consistent patterning of hemimethylation in which most neighbouring dyads shared the same stranded orientation of hemimethylation. As expected, this patterning was absent for simulated reads with ≥3 hemimethylated dyads, for which each CpG had a 50% chance of being methylated (Fig. 1E). Together with the gradual nature of demethylation, and the considerable increase in hemimethylation with ongoing demethylation, this consistent hemimethylation patterning suggests that methylation loss triggered by MEK inhibition is predominantly passive rather than active.

To gain further insight into the mechanisms underlying demethylation in MEKi/L and GSK3i/L, we conducted quantitative tandem mass tag (TMT) whole proteome analysis and RNA-seq on samples matched to the WGBS samples. Interrogation of the DNA methylation machinery showed that *de novo* methyltransferase enzymes are transcriptionally downregulated across GSK3i/L conditions (which trigger only marginal global demethylation) as well as MEKi/L and 2i/L (which induce extensive hypomethylation). This is in agreement with observations from Sim et al. (2017). In contrast, whilst maintenance methylation machinery *Dnmt1* and *Uhrf1* are transcriptionally upregulated across conditions, UHRF1 protein abundance is specifically reduced in 2i/L and MEKi/L (Fig. 1F). Titration of MEK inhibitor concentration within MEKi/L also titrates the level of UHRF1 protein (but not stabilising cofactor USP7), and UHRF1 protein diminishes gradually with duration of MEKi/L culture (Felle et al. 2011) (Fig. 1G & S1F). Taken together, these results suggest that global hypomethylation in 2i/L is driven by MEK inhibition dose-dependent reduction of UHRF1 protein, resulting in gradual, passive loss of DNA methylation via hemimethylation intermediates.

### GSK3i/L-driven impairment of *de novo* methylation results in hypomethylation of enhancers and non-CGI promoters

RT-qPCR analysis of cells cultured in the different conditions for only 24 hours revealed that both GSK3i/L and MEKi/L trigger rapid transcriptional downregulation of the three *de novo* methyltransferases expressed in mESCs, but N2B27 media with LIF alone (N2B27/L) does not. Treatment with an ERK-specific inhibitor in the same basal media with LIF (ERKi/L) resulted in downregulation of *Dnmt3a* and *Dnmt3l* but not *Dnmt3b* (Fig. 2A). Titration of the GSK3 inhibitor within GSK3i/L demonstrated that the resulting transcriptional silencing is dependent upon the dose of inhibitor, most notably for *Dnmt3b* and *Dnmt3l* (Fig. 2B). *Prdm14*, which has been shown to play a role in the transcriptional silencing of *de novo* methyltransferases in 2i/L, was upregulated most significantly in MEKi/L and ERKi/L, but was also more upregulated with increased concentration of GSK3 inhibitor within GSK3i/L up to 3µM (Ficz et al., 2013; Hackett et al., 2013; Leitch et al., 2013; Yamaji et al., 2013) (Fig. 2A & 2B).

**Figure 2:**
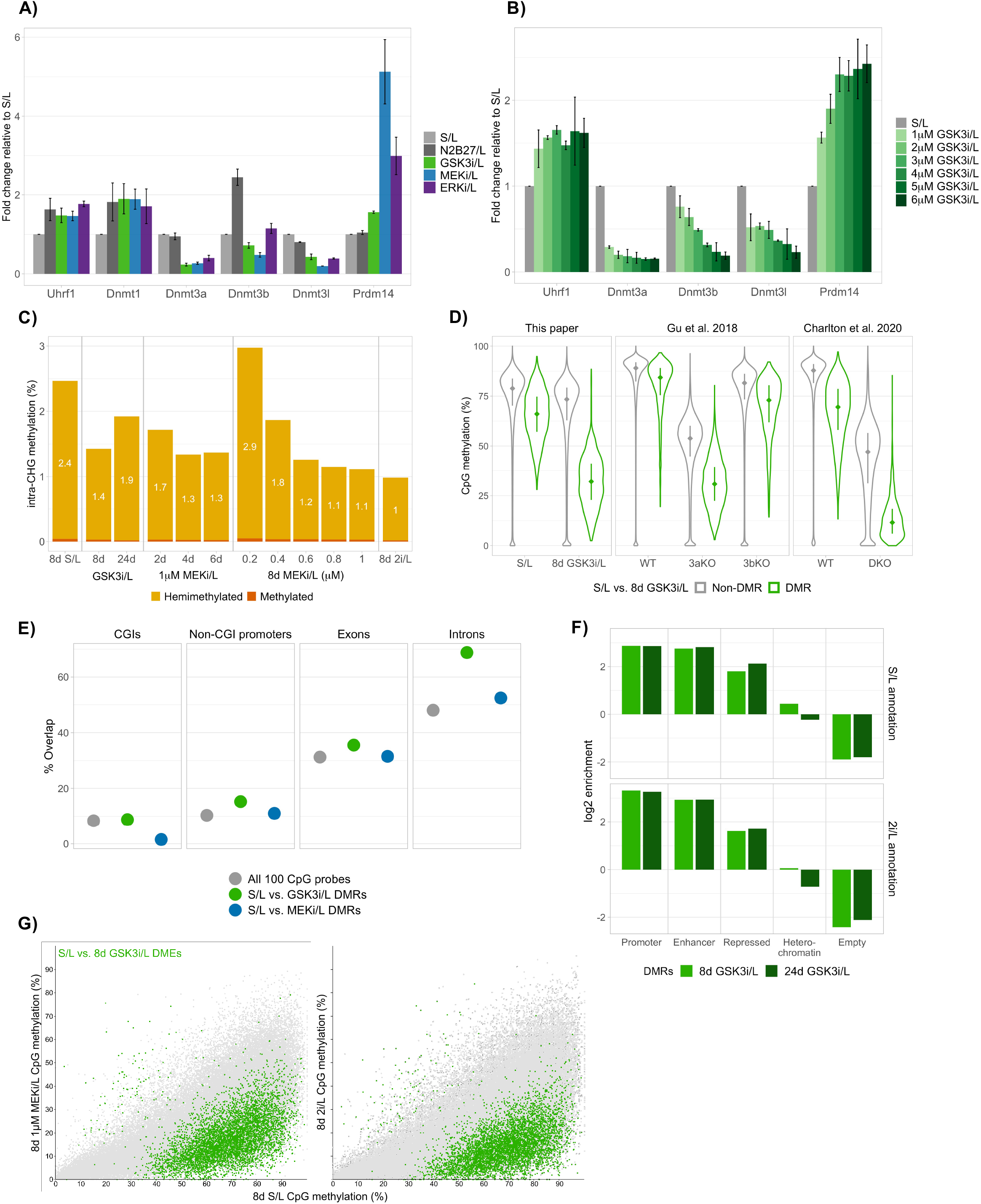
GSK3i/L-driven impairment of *de novo* methylation results in hypomethylation of enhancers and non-CGI promoters. **A)** RT-qPCR analysis of DNA methylation machinery across mESCs cultured for 24h in S/L, N2B27/L, GSK3i/L, MEKi/L or ERKi/L. Error bars show standard deviation. **B)**RT-qPCR analysis of DNA methylation machinery across mESCs cultured for 8 days in GSK3i/L containing different concentrations of GSK3 inhibitor. Error bars show standard deviation. **C)**Percentages of hemimethylated and fully methylated cytosines in a CHG context across culture conditions, calculated using *in silico* strand annealing. **D)** Violin plots depicting percentage CpG methylation levels of S/L vs. 8d GSK3i/L non-DMRs (420,042) and DMRs (2,141), shown across S/L and 8d GSK3i/L samples, Dnmt3a and Dnmt3b single knockout mESCs, and Dnmt3a/3b double knockout mESCs (DKO). **E)** Dotplot depicting percentages of 100 CpG probes that overlap CGIs, non-CGI promoters, exons and introns, shown for all probes, S/L vs. GSK3i/L DMRs and S/L vs. MEKi/L DMRs. **F)** log2 enrichments of S/L vs. 8d GSK3i/L and S/L vs. 24d GSK3i/L DMRs for different classes of genomic element as defined by EpiCSeg (Epigenome Count-based Segmentation) based on S/L and 2i/L cultured mESCs. **G)** S/L vs. 8d GSK3i/L differentially methylated EpiCSeg enhancers (DMEs) highlighted on scatter plots of EpiCSeg enhancer methylation in MEKi/L and 2i/L cultured mESCs versus S/L.

We additionally used *in silico* strand annealing to investigate the levels of non-CpG hemimethylation across culture conditions, considering them a proxy for *de novo* methylation activity (Xu and Corces, 2019). mESCs maintained in S/L exhibited 2.4% non-CpG hemimethylation, those maintained 8 days in 0.2µM MEKi/L 2.9%. However, both cells cultured for longer or in higher concentrations of MEK inhibitor, as well as those cultured in GSK3i/L, showed reduced levels of non-CpG hemimethylation, with 2i/L cultured cells presenting the lowest (Fig. 2C). Taken together with the RT-qPCR data, these results suggest that *de novo* methylation is rapidly suppressed by both MEK and GSK3 inhibition, in the case of MEK inhibitor by concentrations of 0.4µM and above. CpG methylation levels of regions containing hemimethylated CHG sites did not differ from regions containing random CHG sites, suggesting a lack of CpG-methylation-based targeting of CHG methylation (Fig. S2A).

Since mESCs cultured in MEKi/L or 2i/L have both impaired maintenance and *de novo* methylation activity, whilst those cultured in GSK3i/L have reduced *de novo* methylation only, we further investigated methylation changes in GSK3i/L to interrogate specific effects of impaired *de novo* methylation. Differentially methylated 100 CpG regions (DMRs, always compared to S/L samples) in 8d GSK3i/L cultured mESCs were amongst the most hypomethylated regions in Dnmt3a/b double knockout mESCs, confirming a principal role of DNMT3A/B in methylating these regions in S/L (Fig. 2D). However, these DMRs were more hypomethylated in Dnmt3a/b double knockout mESCs than mESCs cultured in GSK3i/L, implying that GSK3i/L causes a partial rather than total impairment of *de novo* methylation activity. Hypomethylation of 8d GSK3i/L DMRs was more pronounced in Dnmt3a single knockout mESCs compared to Dnmt3b knockouts, suggesting that the majority of demethylation in GSK3i/L comes from reduced DNMT3A activity. However, further hypomethylation of these DMRs in the double knockout mESCs points to an additive effect of DNMT3B impairment. Interestingly, 24d GSK3i/L DMRs showed a very similar methylation profile across *de novo* methyltransferase knockouts (Fig. S2B). This suggests that there is no further loss of methylation with prolonged impairment of *de novo* methylation in this system, in contrast to what was originally observed in Dnmt3a/b knockout mESCs (Chen et al., 2003).

Characterisation of GSK3i/L DMRs can therefore provide insight into the genomic regions that are most susceptible to DNA methylation changes when *de novo* methylation activity is modulated. Based on genomic location alone, we found these regions to be strongly enriched for introns, and marginally enriched for exons and non-CGI promoters compared to MEKi/L DMRs (Fig. 2E). In agreement with the enrichment of genic regions, GSK3i/L DMRs were found to be GC rich relative to MEKi/L DMRs and to the genome average, even when regions overlapping promoters and CGIs were excluded (Fig. S2C).

We compared GSK3i/L DMRs to a published EpiCSeg (Epigenome Count-based Segmentation) genome annotation based on 22 ChIP-seq datasets from S/L and 2i/L cultured mESCs (Peng et al., 2020). The DMRs were shown to be enriched for promoters, enhancers and, to a lesser extent, ‘repressed’ regions, which are predominantly characterised by polycomb-deposited marks. This enrichment was very similar for genome segmentations based on either S/L or 2i/L histone marks (Fig. 2F). We henceforth used the enhancer annotations from this EpiCSeg for further analysis. In agreement with the similar enrichment results using S/L or 2i/L EpiCSeg annotations, in GSK3i/L cultured mESCs 2i/L-specific enhancers were only marginally more hypomethylated than S/L-specific enhancers or shared enhancers (Fig. S2D). This suggests that rather than GSK3 inhibition driving demethylation of 2i/L-specific enhancers only, it also triggers some rewiring of enhancers pre-existing in S/L.

Differentially methylated EpiCSeg enhancers (DMEs, always compared to S/L) for 8d GSK3i/L samples were also enriched for introns compared to all enhancers or MEKi/L DMEs (Fig. S2E). EpiCSeg enhancers were found to be GC rich on average compared to 100 CpG probes, whilst GSK3i/L DMEs were no more GC rich (Fig. S2C). This suggests that the elevated GC content of GSK3i/L 100 CpG DMRs can be attributed to an enrichment of these enhancer regions.

We further investigated EpiCSeg enrichments for DMRs of each time point during the MEKi/L culture time course. The earliest time points, at which *de novo* methyltransferases are already transcriptionally silenced but UHRF1 protein is not significantly depleted, were similarly enriched for promoter, enhancer and, to a greater extent, repressed regions. However, as expected, these enrichments diminished during the time course as the loss of DNA methylation became more global. 2i/L DMRs, on the other hand, were marginally more enriched for promoters and enhancers than MEKi/L DMRs, pointing to an effect of GSK3 inhibition within 2i/L. The only notable discrepancy between EpiCSeg enrichments based on S/L or 2i/L genome segmentations was for the earliest time point of MEKi/L culture, for which DMRs were more highly enriched for 2i/L promoters than S/L promoters (Fig. S2F).

DMEs in GSK3i/L cultured mESCs, in which only *de novo* methylation was impaired, were also amongst the most hypomethylated enhancers in MEKi/L and 2i/L, in which both *de novo* and maintenance methylation was impaired (Fig. 2G). This suggests that these impairments compound, resulting, across GSK3i/L, MEKi/L and 2i/L, in most pronounced hypomethylation at regions that are targeted by *de novo* methylation in S/L.

### GSK3i/L hypomethylated enhancers are enriched for 2i/L-specific pluripotency factor binding, TET2 and H3K4me1

Interrogation of EpicSeg enhancers overlapping annotated tissue-specific enhancers (Shen et al., 2012) revealed that only those overlapping mESC enhancers show pronounced hypomethylation in GSK3i/L compared to S/L. Similar analysis for three classes of enhancer defined by Peng et al. – C1 enhancers with STARR-seq activity and active chromatin marks, C2 with STARR-seq activity only, and C3 with active chromatin marks only – showed that C1 and C3 enhancers alone are hypomethylated in GSK3i/L (Peng et al., 2020). C2 enhancers, which may be enhancers that are active in other tissue types, did not show substantial hypomethylation (Fig. S3A).

We proceeded to functionally characterise the enhancers targeted by *de novo* methylation in S/L and differentially methylated in GSK3i/L. These DMEs were enriched for TCF/LEF, KLF and SOX binding motifs compared to a background of non-differentially methylated enhancers (non-DMEs), including for a composite motif of OCT4-SOX2-TCF-NANOG (Fig. 3A). Analysis of MEKi/L DMEs showed no such motif enrichment (Fig. S3B). Use of the Cistrome DB Toolkit (Zheng et al., 2019), which quantifies the overlap between ChIP-seq datasets in the database and genomic regions of interest, showed the GSK3i/L DMEs to be most similar to binding profiles of CTNNB1 (β-CATENIN, the downstream effector of GSK3 inhibition), TFCP2L1, TCF3, EP300, POU5F1 (OCT4) and SOX2. In the case of CTNNB1, Cistrome datasets of S/L mESCs cultured with GSK3 inhibition were much more similar to DMEs than those of S/L mESCs without GSK3 inhibition. For NANOG, EP300 and OCT4, the datasets most similar to DMEs were those of cells grown in 2i/L. Amongst histone mark datasets, H3K4me1 profiles were most similar to GSK3i/L DMEs, again principally datasets from mESCs cultured in 2i/L. A size-matched set of non-DMEs did not show as significant an overlap with any transcription factor (TF) binding profiles in the Cistrome database, and did not share any top enrichments with those of DMEs. Non-DMEs showed elevated similarity to H3K27ac profiles, but not H3K4me1, whilst DMEs did not show high similarity to H3K27ac distribution (Fig. S3C).

**Figure 3:**
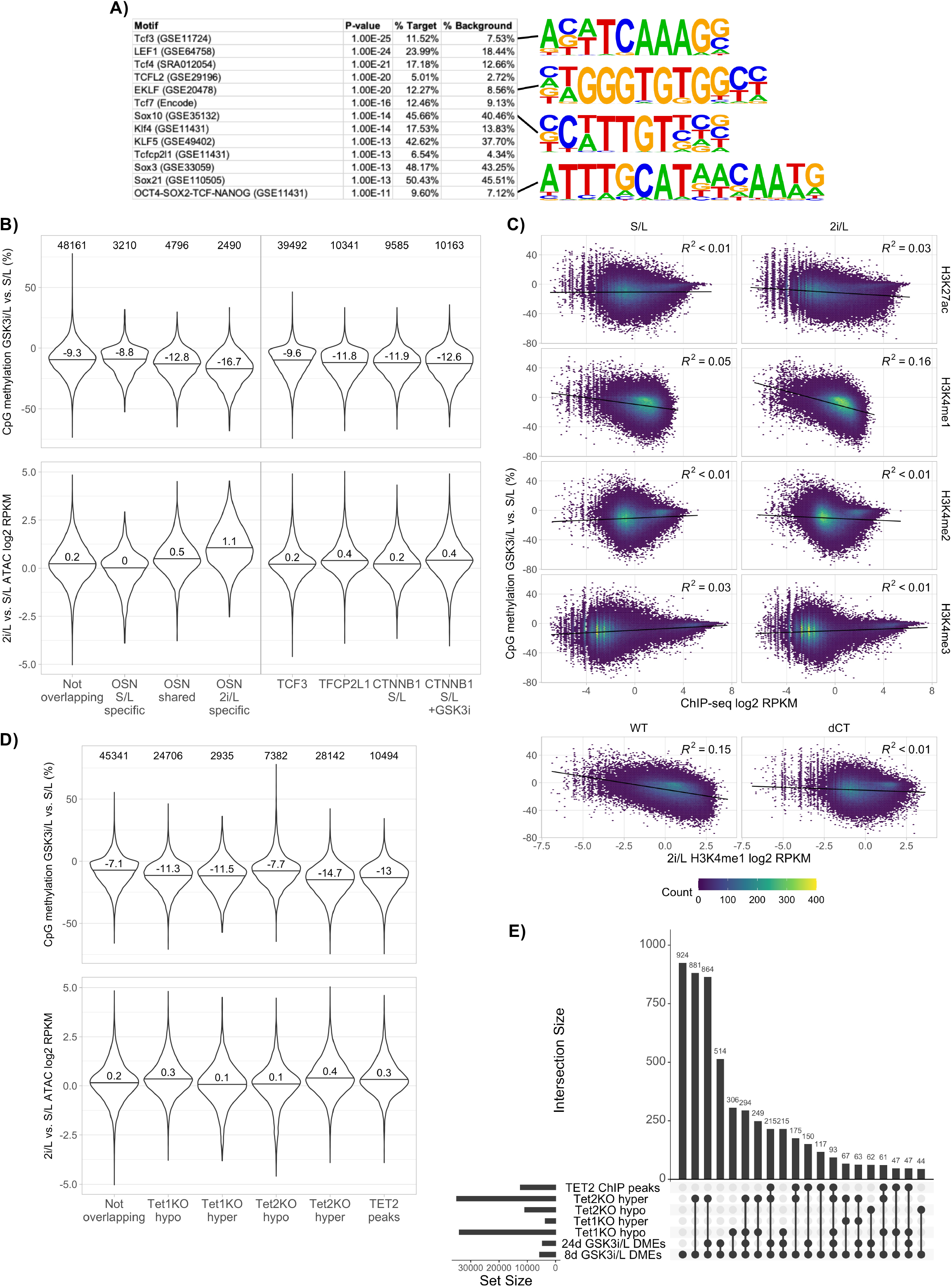
GSK3i/L hypomethylated enhancers are enriched for 2i/L-specific pluripotency factor binding, TET2 and H3K4me1. **A)** The top results of Homer motif analysis for differentially methylated enhancers (DMEs) between S/L and 8d GSK3i/L. **B)**Upper - violin plots depicting percentage CpG methylation change between S/L and 8d GSK3i/L at EpiCSeg enhancers overlapping selected transcription factor peaks. Lower – ATAC-seq signal in 2i/L versus S/L at the same enhancer classes. The number of enhancers in each class is printed above the violins. **C)**Scatter plots coloured by density showing the relationship between percentage CpG methylation change between S/L and 8d GSK3i/L and abundance of different histone marks in S/L or 2i/L at EpiCSeg enhancers. The lower panel shows just 2i/L H3K4me1 in wildtype (WT) mESCs or mESCs catalytically deficient for both MLL3 and MLL4 (dCT). Across all panels, a small number of outliers with methylation change >70% or ChIP-seq log2 RPKM < - 7 were not plotted purely to aid visualisation. **D)** Upper - violin plots depicting percentage CpG methylation change between S/L and 8d GSK3i/L at EpiCSeg enhancers overlapping DMRs in Tet single knockout mESCs or 2i/L TET2 ChIP-seq peaks. Lower – ATAC-seq signal in 2i/L versus S/L at the same enhancer classes. The number of enhancers in each class is printed above the violins. **E)** Upset plot illustrating the degree of overlap between 8d and 24d GSK3i/L DMEs and DMRs in Tet single knockout mESCs or 2i/L TET2 ChIP-seq peaks. The plot is 8d GSK3i/L DME centric, i.e. shows the top largest overlaps involving 8d GSK3i/L DMEs. Each overlap quantified is a distinct overlap, meaning the regions in that overlap do not also overlap with any other set of regions amongst those analysed.

To validate these (closely-aligned) transcription factor motif and binding profile enrichments, we used publicly available ChIP-seq datasets to examine DNA methylation at enhancers overlapping peaks of TF binding. The most significant hypomethylation was observed at enhancers harbouring OCT4, SOX2 and NANOG (OSN) binding specific to 2i/L cultured mESCs, whilst enhancers with S/L-specific OSN binding were no more hypomethylated than those without peaks of any of the TFs investigated (Fig. 3B). This remained the case when OCT4, SOX2 and NANOG bound enhancers were analysed separately, with enhancers that have 2i/L-specific SOX2 enrichment the most hypomethylated of all (Fig. S3E). TCF3 peaks are extremely abundant in mESCs, and therefore, unsurprisingly, the largest overlap between DMEs and TF peaks amongst factors we examined was for TCF3 (Fig. S3D). In line with their abundance, however, enhancers overlapping TCF3 peaks did not show substantial hypomethylation compared to non-overlapping enhancers. Enhancers with TFCP2L1 or CTNNB1 binding were only marginally hypomethylated compared to non-overlapping enhancers (Fig. 3B). Interestingly, the average difference in ATAC-seq signal between 2i/L and S/L cultured mESCs anticorrelated with DNA methylation change over the different enhancer sets, suggesting that enhancers that are hypomethylated in GSK3i/L (which are likewise amongst the most hypomethylated in 2i/L – see Fig. 2G) also become more accessible in 2i/L. 2i/L-specific OSN-bound enhancers had the most accessible chromatin in 2i/L compared to S/L, and indeed GSK3i/L DMEs became more accessible than non-DMEs (Fig. 3B & S3F).

To interrogate the implication from Cistrome Toolkit histone mark analysis that DMEs overlap most significantly with H3K4me1, we analysed the correlation at enhancers between DNA methylation change in GSK3i/L versus S/L, and the abundance of different histone marks in S/L and 2i/L. No correlation was observed between DNA methylation change and H3K27ac, H3K4me2 or H3K4me3 enrichment in either S/L or 2i/L. However, whilst there was little correlation with H3K4me1 of mESCs cultured in S/L, a negative correlation was observed for mESCs in 2i/L (Fig. 3C). This correlation was weak, but markedly increased compared to all other comparisons. This same correlation was evident in an independent dataset, but completely lost in cells with catalytically dead MLL3 and MLL4, the two main writers of H3K4me1 in mESCs (Fig. 3C). Therefore, the more H3K4me1 added *de novo* to an enhancer in 2i/L, the more hypomethylated that enhancer is likely to be in GSK3i/L. The correlations between H3K4me1 and DNA methylation loss in MEKi/L and 2i/L, in which maintenance methylation is also impaired, are markedly reduced (Fig S3G).

Previous reports have demonstrated a role for active demethylation by TET proteins in the loss of DNA methylation observed when *de novo* methyltransferases are ablated, as well as a particular role of TET2 in promoting naïve pluripotent cell identity (Charlton et al. 2020; Fidalgo et al. 2016; Gu et al. 2018; Jeong et al. 2014). We therefore analysed the DNA methylation in GSK3i/L relative to S/L of enhancers overlapping DMRs in Tet1 and Tet2 single knockout mESCs. Both 8d and 24d GSK3i/L DMEs overlapped most frequently with sites that are hypermethylated in Tet2 knockout cells (Fig. 3E). Enhancers overlapping these Tet2 knockout hypermethylated regions, as well as those overlapping 2i/L TET2 ChIP-seq peaks, were the most hypomethylated in GSK3i/L, suggesting a link to TET2 catalytic activity. Again, these were also the enhancers with the highest chromatin accessibility in 2i/L compared to S/L (Fig. 3D).

Furthermore, TET2 binding in 2i/L was more enriched at both 8d and 24d GSK3i/L DMEs compared to non-DMEs (Fig S3H). This suggests that, upon impairment of *de novo* methylation in GSK3i/L, the balance between *de novo* methylation and active demethylation by TET2 at enhancers is tipped, and enhancers subject to demethylation by TET2 are amongst the most hypomethylated.

### Enhancer hypomethylation coincides with upregulation of proximal genes

The gene expression signatures of mESCs cultured for 8 days in either GSK3i/L or MEKi/L both lie between those of S/L and 2i/L cultured cells according to PC1 of RNA-seq principal component analysis, which captures 47% of variation in the data (Fig. 4A). Accordingly, genes that are upregulated or downregulated in 2i/L compared to S/L are also up or downregulated, respectively, in GSK3i/L and MEKi/L, but to a lesser extent. And genes up or downregulated in GSK3i/L are further up- or downregulated, respectively, in 2i/L (Fig. 4B).

**Figure 4:**
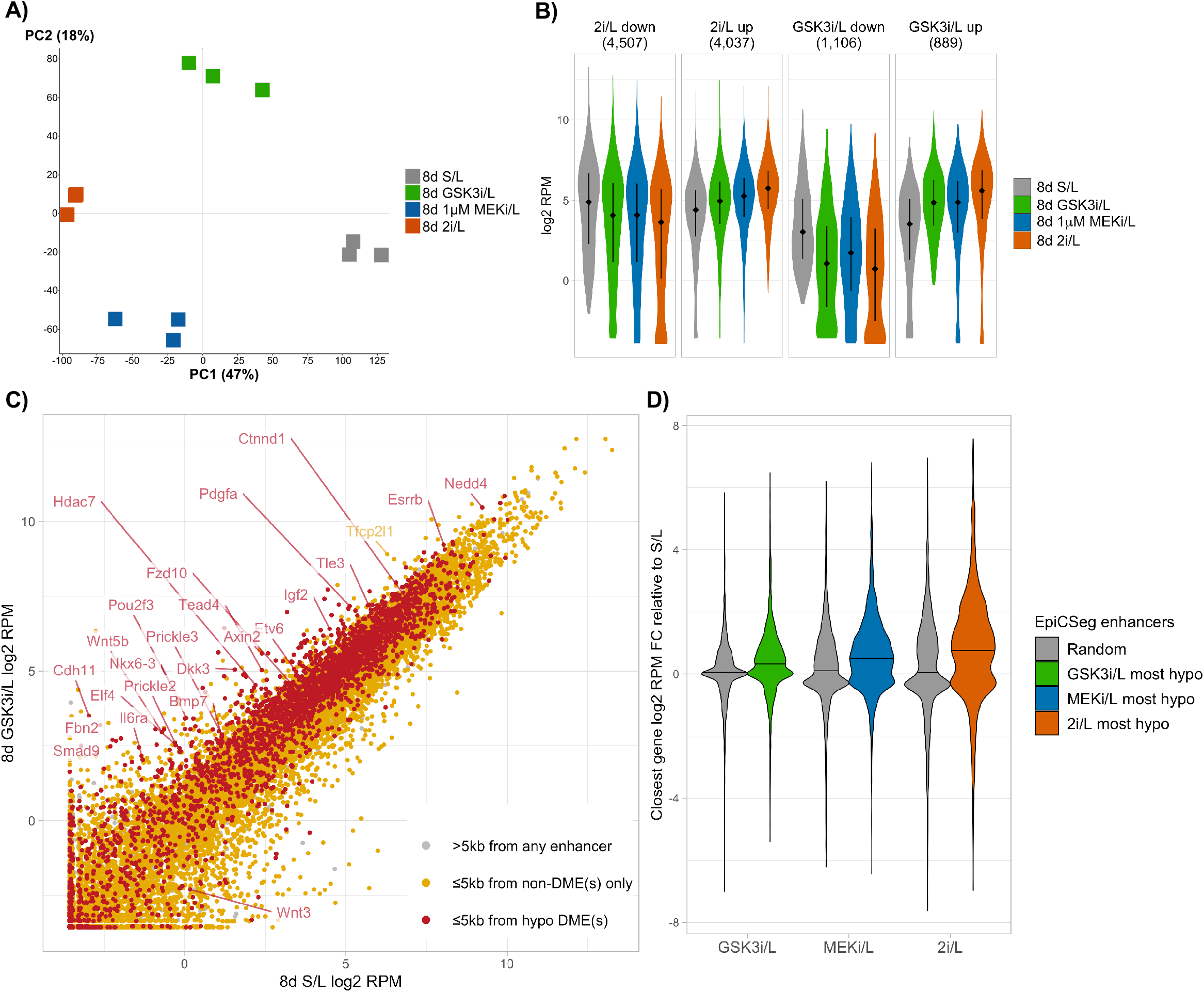
Enhancer hypomethylation coincides with upregulation of proximal genes. **A)** Principal component analysis of biological triplicate RNA-seq samples for 8-day culture of mESCs in S/L, 2i/L, GSK3i/L or 1µM MEKi/L. **B)**Violin plots depicting gene expression (log2-transformed RNA-seq reads per million [RPM]) of genes that are either downregulated or upregulated in either 2i/L or GSK3i/L compared to S/L, across samples. Genes were called as differentially expressed according to a DESeq test with p<0.01. The number of differentially expressed genes in each group is printed above each panel. **C)**Scatter plot showing gene expression (log2 RPM) in S/L versus 8d GSK3i/L cultured mESCs. Datapoints for genes >5kb from any enhancer are coloured grey, those for genes ≤5kb from non-DMEs only are coloured yellow, and those for genes ≤5kb from an enhancer hypomethylated in GSK3i/L are coloured red. Datapoints are plotted in layers in the same order, with those for genes >5kb from any enhancer plotted beneath those for genes ≤5kb from non-DMEs, which are beneath those for genes ≤5kb from a hypomethylated enhancer. **D)** Violin plots depicting gene expression (log2 RPM) fold change between S/L and GSK3i/L, MEKi/L or 2i/L. Expression is plotted of genes that are the closest within 100kb to either random enhancers, or the 5,000 most hypomethylated enhancers in GSK3i/L, MEKi/L or 2i/L, compared to S/L. These 5,000 enhancers were selected having pre-excluded enhancers overlapping -5000bp to +500bp promoter regions. The enhancer methylation comparisons correspond to the gene expression comparisons, i.e. closest gene expression in GSK3i/L compared to S/L is plotted for the 5,000 most hypomethylated enhancers in GSK3i/L compared to S/L. Random enhancers are 5,000 randomly selected enhancers (not overlapping promoters) that are not amongst the 5,000 most hypomethylated in GSK3i/L, MEKi/L or 2i/L.

To investigate whether loss of DNA methylation at enhancers in GSK3i/L is associated with changes in enhancer function, we analysed expression of genes within 5 kilobases (kb) of enhancers. Genes within 5kb of one or more hypomethylated enhancers exhibited higher gene expression in GSK3i/L relative to S/L compared to genes within 5kb of non-DMEs only, as well as compared to genes more than 5kb from any enhancer (Fig. 4C).

We subsequently analysed the 5,000 most hypomethylated enhancers (disregarding any that overlapped promoter regions) in each of GSK3i/L, MEKi/L and 2i/L compared to S/L, which were on average approximately 30% hypomethylated in GSK3i/L and 60% hypomethylated in MEKi/L and 2i/L compared to S/L (Fig. S4A). As our previous analysis (Fig. 2G) suggested, there was considerable overlap between the most hypomethylated enhancers across conditions, with over a fifth shared across all three culture conditions (Fig. S4C). We interrogated the expression of the closest gene within 100kb to each of these enhancers, as well as the closest gene within 100kb to a random set of 5,000 enhancers that were not amongst the most hypomethylated in any condition. Consistently across conditions, relative to S/L, the closest genes to the most hypomethylated enhancers were upregulated compared to genes closest to random enhancers (Fig. 4D). In line with the overlap between the most hypomethylated enhancers across conditions, there was also substantial overlap in the closest genes to these (Fig. S4D). Perhaps as a result of this, the degree of upregulation of enhancer-proximal genes across these sample comparisons mirrored the trend of genes upregulated in 2i/L, with more upregulation in 2i/L compared to in GSK3i/L or MEKi/L (Fig. 4B & 4D).

To assess whether this connection between enhancer hypomethylation and gene upregulation is unique to proximal genes, and to analyse this relationship for genes for which a promoter-enhancer interaction is known in mESCs, we additionally interrogated the expression of genes linked by Capture Hi-C (CHi-C) to the 5,000 most hypomethylated enhancers across GSK3i/L, MEKi/L and 2i/L. These genes showed the same trend as those proximal to hypomethylated enhancers, although genes CHi-C linked to random enhancers were also slightly upregulated across culture conditions, and therefore upregulation relative to these was more marginal (Fig. S4B). Again, as expected based on the overlap between the most hypomethylated enhancers across conditions, there was also considerable overlap in the genes CHi-C linked to these enhancers (Fig. S4E).

The majority of genes close or CHi-C linked to the most hypomethylated enhancers in 2i/L also showed increased expression at the protein level, demonstrating that elevated gene expression coincident with enhancer hypomethylation has a functional consequence in naïve pluripotent mESCs (Fig. S4F).

## Discussion

For over a decade, mouse embryonic stem cells have been cultured with inhibitors of two key signalling pathways in the absence of a comprehensive understanding of the individual effects these have upon the epigenome. Here we demonstrate that both MEK and GSK3 inhibition trigger significant, but distinct, changes in the DNA methylome of mESCs. Whilst MEK inhibition triggers both rapid impairment of *de novo* methylation and a slower, dose-dependent impairment of maintenance methylation, GSK3 inhibition causes impairment of *de novo* methylation alone. This modulation of *de novo* methylation resulted in hypomethylation at enhancers, and particularly those with abundant pluripotency factor, TET2 and H3K4me1 binding specifically in 2i/L (Fig. 5).

**Figure 5:**
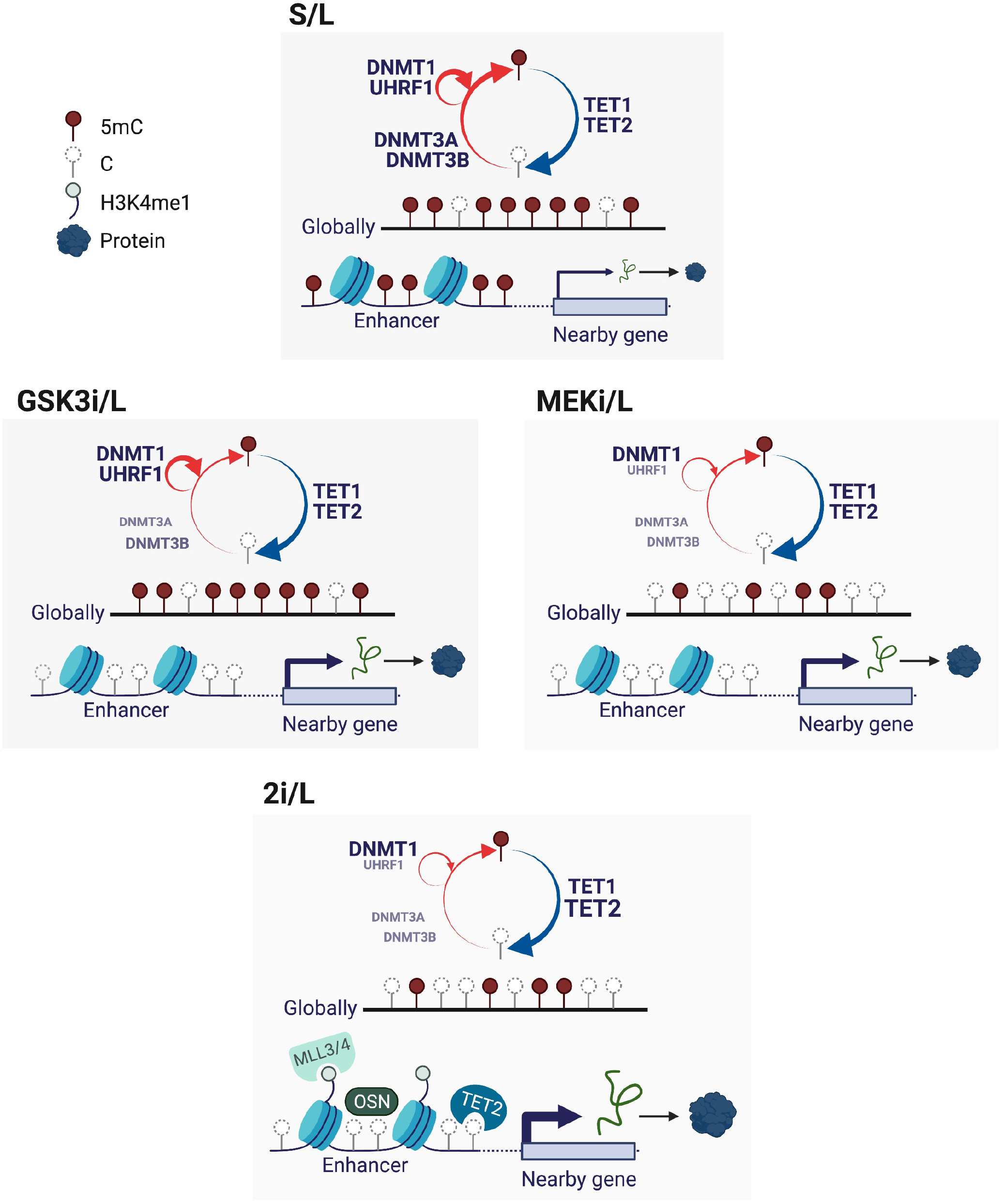
The same example enhancer is shown across culture conditions. In S/L, all three de novo methyltransferases expressed in mESCs are abundant, as is the maintenance methylation machinery DNMT1 and UHRF1, as well as DNA demethylating TET1/2. Genome-wide DNA methylation levels are high, but there is constant turnover at regions targeted by both DNMT3 and TET enzymes (Rulands et al. 2018; Gu et al. 2018). In GSK3i/L, all three de novo methyltransferases are rapidly transcriptionally silenced whilst maintenance methylation machinery persists. Genome-wide DNA methylation levels remain high, but there is targeted hypomethylation at enhancers, particularly those that have MLL3/4-deposited H3K4me1, OCT4, SOX2 and NANOG (OSN) binding, and TET2 enrichment in 2i/L. In MEKi/L and 2i/L, UHRF1 protein is also depleted, and the genome becomes globally hypomethylated. Across GSK3i/L, MEKi/L and 2i/L there is substantial overlap in the enhancers that are most hypomethylated, and genes proximal to these enhancers are upregulated both transcriptionally and, in the majority of cases, also at the protein level. Figure created with BioRender.com.

The impairment of both *de novo* and maintenance methylation by MEK inhibition has an additive effect, so that enhancers targeted by *de novo* methylation in S/L that are hypomethylated in GSK3i/L are also amongst the most hypomethylated in MEKi/L and 2i/L. There is considerable overlap across these conditions, therefore, in the genes proximal or CHi-C linked to the most hypomethylated enhancers, which are transcriptionally upregulated in response to both GSK and MEK inhibition.

In naïve pluripotent mESCs, and not dissimilarly in naïve human ESCs, MEK inhibition impedes differentiation signals, whilst GSK3 inhibition promotes clonogenicity and self-renewal under these conditions (Takashima et al., 2014; Ying et al., 2008). In the pre-implantation mouse embryo, inhibition of MEK results in an expansion of pluripotent epiblast and reduction of hypoblast and, again, GSK3 inhibition appears to support the expansion of pluripotent cells under certain conditions (Nichols et al., 2009). Canonical Wnt signalling, driven by GSK3 inhibition, causes CTNNB1 (β-CATENIN) to enter the nucleus and bind alongside TCF3 at regulatory regions both with and without pluripotency factor binding, which regulate genes related to pluripotency or lineage specification, respectively (Zhang et al., 2013). Although we find enrichment of the TCF3 binding motif and CTNNB1 binding within GSK3i/L DMEs, hypomethylation is more pronounced at enhancers with binding of OCT4, SOX2 and NANOG specifically in 2i/L, suggesting a stronger link between DNA methylation loss and transcription factor binding at enhancers related to naïve pluripotency.

Our observation that DMRs in GSK3i/L are more hypomethylated in Dnmt3a than Dnmt3b single knockout mESCs is in agreement not only with the global methylation levels of these knockouts, but also with the observation that DNMT3A is the principal methyltransferase at pluripotency gene enhancers (Petell et al., 2016). All DNMT3 enzymes harbour an ATRX-DNMT3L-DNMT3A (ADD) domain, which binds unmethylated H3K4 but is repelled by additional H3K4 methyl moieties (Ooi et al., 2007; Zhang et al., 2010). CpG-rich promoters with high levels of H3K4me3 are consistently lowly methylated, but enhancers marked with H3K4me1 exhibit more intermediate and heterogeneous levels of DNA methylation in S/L-cultured mESCs (Rulands et al., 2018; Sharifi-Zarchi et al., 2017). Our data suggests that when *de novo* methyltransferases are downregulated in response to GSK3 inhibition, the reduced quantities of enzyme are no longer able to methylate regions marked by MLL3/4-deposited H3K4me1, and therefore these are amongst the most hypomethylated regions. Alternatively, or perhaps additionally, deposition of H3K4me1 by MLL3/4 may be facilitated by loss of DNA methylation.

An outstanding and compelling question following our investigation is that of the temporal and mechanistic hierarchy of changes in DNA methylation, H3K4 mono-methylation, and pluripotency factor binding upon GSK3 inhibition. Given the absence of genetic manipulation, and the rapidity with which GSK3 inhibition triggers downregulation of all three *de novo* methyltransferases in mESCs, culture in GSK3i/L provides a valuable system with which to investigate such mechanistic questions. Our study lays key groundwork for this by providing detailed insights into the epigenetic, and associated transcriptional, impacts of inhibiting GSK3 signalling in mouse pluripotent cells.

## Methods

### Cell culture

E14Tg2a (XY) mouse embryonic stem cells (mESCs) were cultured feeder-free in normoxia on 0.1% gelatin-coated plates. 1.5 million cells were plated per 10cm plate in S/L media (DMEM supplemented with 15% fetal bovine serum, 1% Penicillin-Streptomycin, 1% MEM non-essential amino acids, 1% Glutamax, 50 μM *β*-mercaptoethanol [all Gibco] and 10^3^ U LIF [Cambridge Stem Cell Institute]). The following day, cells were washed twice with PBS and the media changed to S/L, 2i/L, GSK3i/L or MEKi/L media. 2i/L, GSK3i/L and MEKi/L media had in common the basal N2B27 media with LIF (1:1 ratio DMEM/F12 to neurobasal medium, 0.5% 100X N-2 supplement, 1% 50X B-27 supplement without Vitamin A, 1% Glutamax, 1% Penicillin-Streptomycin, 50 μM *β*-mercaptoethanol [all Gibco] and 10^3^ U LIF). 2i/L media was supplemented with 1μM PD03259010 and 3μM CHIR99021 (both Cambridge Stem Cell Institute), GSK3i/L media with 3μM CHIR99021 only, and MEKi/L with 1μM PD03259010 only. Cells were passaged every 2 days using TrypLE Express (Gibco), and 3 million cells were plated per 10cm plate.

For collection at designated time points for RNA and DNA extraction, cells were washed twice with cold PBS and then scraped from plates in RLT plus buffer (Qiagen). For collection for whole proteome mass spectrometry, cells were washed twice with cold PBS containing protease and phosphatase inhibitors, pelleted, and flash frozen.

For the 24-hour cell culture experiment, N2B27/L (N2B27 basal media with 10^3^ U LIF) and ERKi/L (N2B27 basal media with 1µM ERK-specific inhibitor SCH772984 and 10^3^ U LIF) media were also used.

### Whole genome bisulphite sequencing (WGBS)

DNA was extracted using the DNA/RNA mini kit (Qiagen) and quantified using a Qubit Fluorometer. DNA was fragmented to an average of 400bp using a Covaris sonicator, and 50ng of sonicated DNA together with a spike-in of 8ng lambda DNA was used for library preparation. Methylated adaptors were added using NEBNext Ultra II DNA Library Prep reagents, then samples were bisulphite converted using the Zymo EZ Methylation-Direct kit. Libraries were amplified using KAPA HiFi Uracil+ then analysed on a Bioanalyzer and by KAPA quantification. A 10nM pool of 26 duplicate samples was 100bp paired-end sequenced over two lanes of a NovaSeq S1 flowcell at the Cancer Research UK Cambridge Institute.

### RNA-sequencing

RNA was extracted from frozen cell pellets using the DNA/RNA mini kit (Qiagen) and subsequently DNase treated using the TURBO DNA-free kit ‘Rigorous DNase treatment’ protocol. Libraries were made by the NGS library facility at the Cambridge Stem Cell Institute, using the NEBNext Poly(A) mRNA Magnetic Isolation Module combined with the NEBNext Ultra II Directional Library Prep Kit for Illumina. Final libraries were KAPA quantified and the library quality assessed with the Agilent Bioanalyser DNA High Sensitivity Kit. A 20nM pool of 17 libraries was sequenced over two lanes of HiSeq2500-RapidRun 50bp Single End sequencing at the Babraham Institute.

### Tandem mass tag (TMT) 16-plex quantitative whole proteome analysis

The mESC pellets were lysed with dissolution buffer (100mM triethylammonium bicarbonate [Sigma, #T4708], 1% Sodium deoxycholate [SDC], 10% isopropanol, 50mM NaCl) followed by tip sonication and boiling at 90°C for 5 minutes. The protein concentration was estimated using the Bradford assay (Bio-Rad, Quick StartTM); 70µg total protein per sample was reduced with tris-2-carboxyethyl phosphine (TCEP, Sigma) for 1 hour at 60°C at a final concentration of 5mM, followed by cysteine blocking for 10 minutes at room temperature using methyl methanethiosulfonate (MMTS, Sigma) at a final concentration of 10mM. Samples were digested overnight at 37°C with trypsin (Pierce #90058) and the next day peptides were labelled with the TMTpro16plex reagents according to manufacturer’s instructions (Thermo Scientific).

After labelling, samples were pooled and 20µl formic acid was added followed by centrifugation at 10,000 rpm for 5 minutes to remove SDC. The TMT mixture was fractionated on a Dionex UltiMate 3000 system at high pH using the X-Bridge C18 column (3.5μm 2.1×150mm, Waters). Fractions were analyzed on a Dionex UltiMate 3000 UHPLC system coupled with the nano-ESI Fusion-Lumos (Thermo Scientific) mass spectrometer. Samples were loaded on the Acclaim PepMap 100, 100μm × 2cm C18, 5μm, 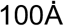 trapping column with the µlPickUp injection method at loading flow rate 5μL/min for 10 minutes. For the peptide separation, the EASY-Spray analytical column 75μm × 25cm, C18, 2μm, 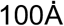 column was used for multi-step gradient elution. The full scans were performed in the Orbitrap in the range of 380-1500 m/z at 120K resolution and the MS2 scans were performed in the ion trap with collision energy 32%. Peptides were isolated in the quadrupole with isolation window 0.7Th. The 10 most intense fragments were selected for Synchronous Precursor Selection (SPS) HCD-MS3 analysis with MS2 isolation window 2.0 and HCD collision energy 50%. The detection was performed with Orbitrap resolution 50K and in scan range 100-400 m/z.

Raw data were processed with the SequestHT search engine on Proteome Discoverer 2.4 software and searched against a Uniprot database containing mouse reviewed entries. The parameters for the SequestHT node were as follows: Precursor Mass Tolerance 20ppm, Fragment Mass Tolerance 0.5Da, Dynamic Modifications were Oxidation of M (+15.995Da), Deamidation of N, Q (+0.984Da) and Static Modifications were TMTpro at any N-Terminus, K (+304.207Da) and Methylthio at C (+45.988Da). The consensus workflow included TMT signal-to-noise (S/N) calculation and the level of confidence for peptide identifications was estimated using the Percolator node with decoy database search. Strict FDR was set at q-value<0.01. For the downstream data analysis, the R package qPLEXanalyzer (Papachristou et al., 2018) was used. Peptide intensities were normalised by median scaling, and then summed to get protein intensities.

The TMT mass spectrometry proteomics data have been deposited to the ProteomeXchange Consortium via the PRIDE partner repository (Perez-Riverol et al., 2019) with the dataset identifier PXD029686. Only S/L replicates 1-3 were used for analysis in this paper, since these were the replicates matched to RNA-seq and WGBS samples.

### Western blotting

Cell pellets were incubated 20 minutes with RIPA lysis buffer (50 mM Tris-HCl pH7.5, 300 mM NaCl, 1 mM EDTA, 1% Triton X-100, 0.1% SDS, 1X Roche COMPLETE Mini EDTA-free protease inhibitor) on ice, then centrifuged and supernatants flash frozen. Protein concentrations were quantified using Bio-Rad Protein Assay Dye Reagent, and 25µg protein together with 1.5% β-mercaptoethanol loading buffer were boiled at 95°C for 5 minutes prior to loading. Samples were run in precast NuPAGE Bis-Tris gels using either NuPAGE MES or MOPS SDS Running Buffer, then transferred onto PVDF membrane (Immobilon-P 0.45 μm, Millipore) either at 25V overnight or at 400mA constant for 90 minutes (both at 4°C and using NuPAGE Transfer Buffer with 20% methanol).

Membranes were blocked in PBS 0.05% Tween (PBST) 5% skimmed milk powder (w/v) (PBST- M) for one hour, then incubated with primary antibodies diluted in PBST-M for two hours.

Following extensive washing with PBST, membranes were incubated with HRP-conjugated secondary antibodies diluted in PBST-M for one hour. p-P90RSK and p-ERK antibodies were diluted in TBS 0.05% Tween (TBST) 5% BSA and membranes also washed with TBST. Membranes were developed using Amersham Western Blotting Detection Reagents.

*Western blot antibodies used in this study (M=monoclonal; P=polyclonal):*

| Antibody | Type | Host | Reference | Dilution |
| --- | --- | --- | --- | --- |
| UHRF1 | 1°, P | Rabbit | Santa Cruz, sc-98817 (M-132) | 1:3000 |
| UHRF1 | 1°, M | Rabbit | Santa Cruz, sc-373750 (H-8) | 1:2000 |
| p-P90RSK (Thr359) | 1°, M | Rabbit | CST, #8753 | 1:1000 |
| p44/42- ERK1/2 | 1°, M | Rabbit | CST, #4370 | 1:1000 |
| USP7 | 1°, M | Mouse | Santa Cruz, sc-137008 | 1:1000 |
| DNMT3B | 1°, M | Mouse | Abcam, ab13604 | 1:600 |
| β-ACTIN | 1°, M | Mouse | Abcam, ab6276 | 1:40,000 |
| Anti-rabbit IgG- HRP | 2°, P | Goat | Santa Cruz, sc-2030 | 1:3000-1:6000 |
| Anti-rabbit IgG- HRP | 2°, P | Goat | Bio-Rad, #1706515 | 1:3000 |
| Anti-mouse IgG-HRP | 2°, P | Goat | Santa Cruz, sc-2005 | Variable |

### Reverse transcription quantitative polymerase chain reaction (RT-qPCR)

RNA was extracted from frozen cell pellets using the RNeasy Mini Kit (Qiagen) and subsequently DNase treated using the TURBO DNA-free kit ‘Rigorous DNase treatment’ protocol. DNAse-treated RNA was reverse transcribed using the RevertAid First Strand Kit with random hexamer primers. The resulting cDNA was diluted 1:50 in a reaction with Platinum™ SYBR Green qPCR SuperMix-UDG (ThermoFisher Scientific) and 1µM primers. The average expression of housekeeping genes Atp5b and Hspcb was used to normalise gene expression relative to S/L samples using the ddCt method.

*RT-qPCR primers used in this study:*

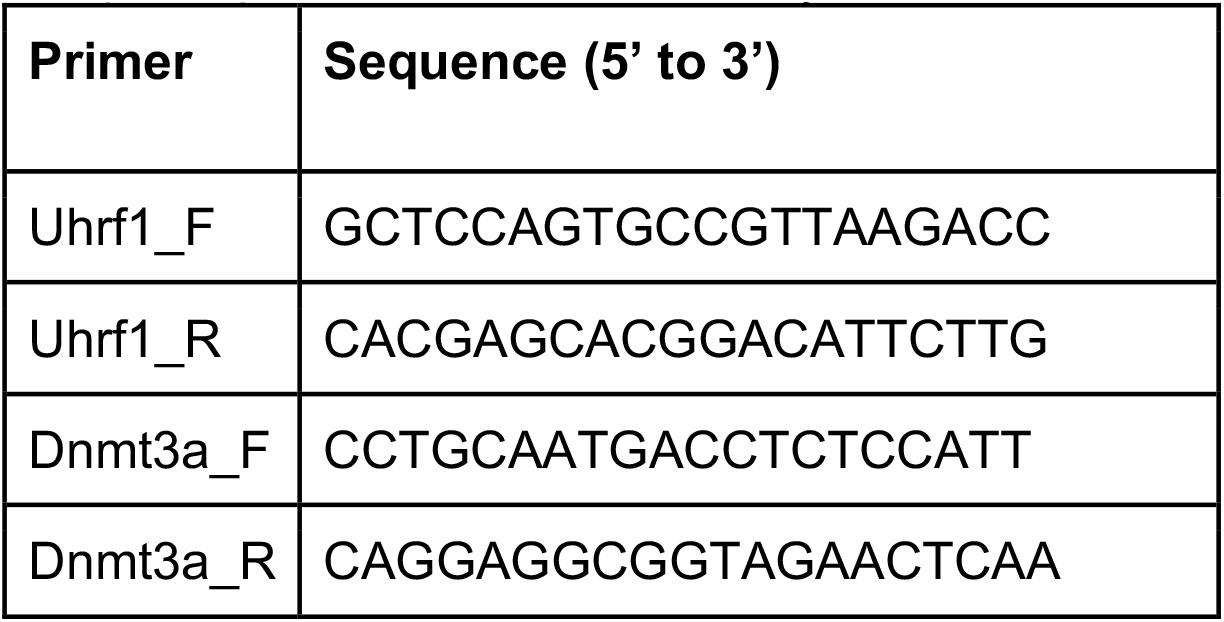
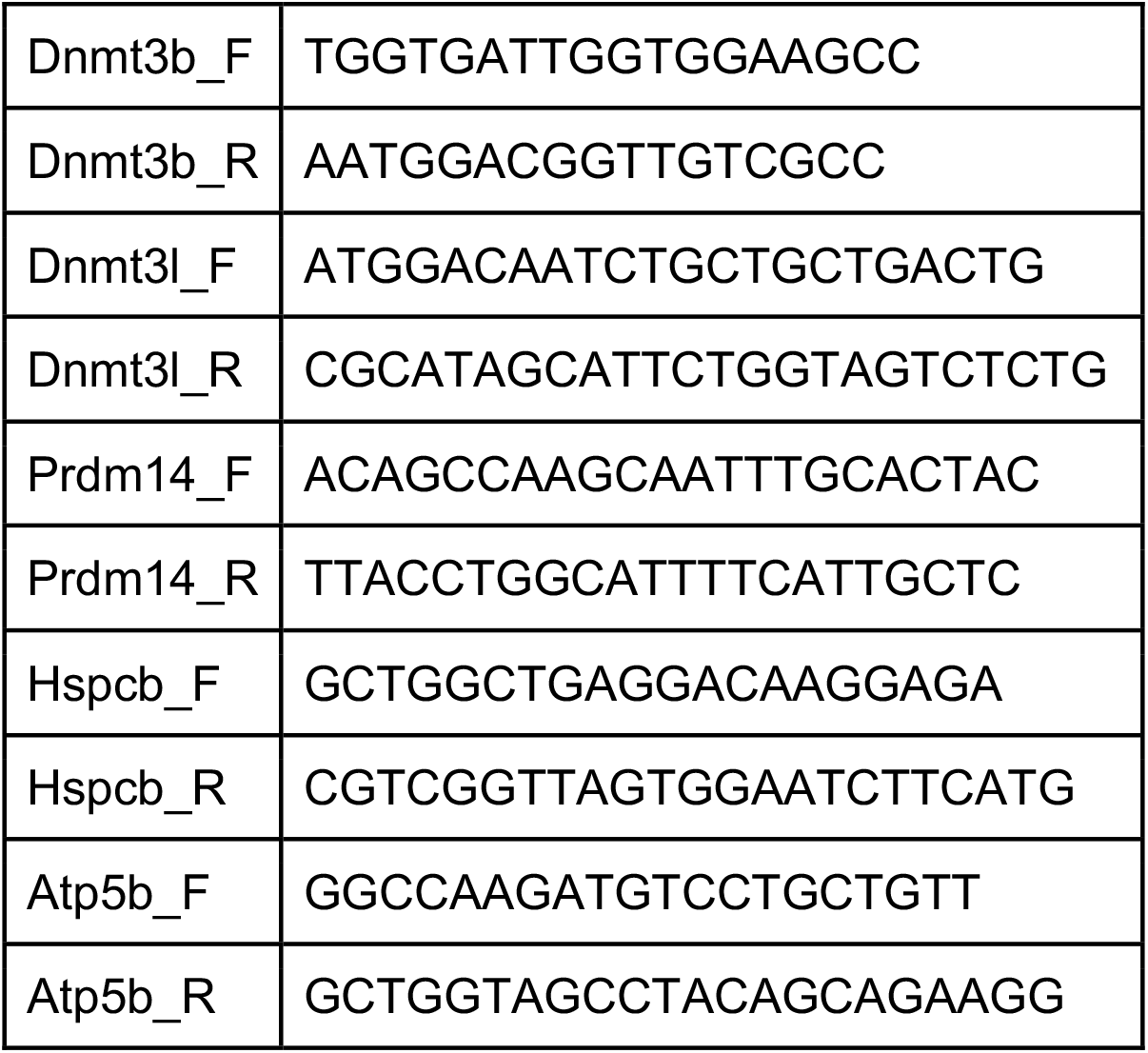

### WGBS analysis

Raw sequence paired-end reads were trimmed to remove both poor quality calls and adapters using Trim Galore (v.0.6.5, https://www.bioinformatics.babraham.ac.uk/projects/trim_galore/). Sequences were mapped to the mouse GRCm38 genome assembly using Bismark (v0.22.3, Krueger and Andrews [2011]), deduplicated with deduplicate_bismark, then CpG methylation calls were extracted and analysed with SeqMonk (https://www.bioinformatics.babraham.ac.uk/projects/seqmonk/). Initial, genome-wide analysis was carried out over 100 CpG probes with at least 20 observations. Repetitive element annotations were generated by RepeatMasker (http://www.repeatmasker.org), sourced via the UCSC Genome Browser (GRCm38, January 2021). DMRs were defined within Seqmonk as 100 CpG probes with minimum 20 observations, which passed both a logistic regression test p≤0.01 and an EdgeR test p≤0.01, and had a minimum methylation difference of 30%.

EpiCSeg-based enrichment analysis was performed using a purpose-built Shiny app (https://www.bioinformatics.babraham.ac.uk/shiny/ChromHMM_state_explorer/) and S/L and 2i/L chromatin segmentations from Peng et al. (2020). S/L and 2i/L enhancer annotations from this EpiCSeg segmentation were taken together for further analysis, with duplicates removed, overlapping enhancers merged, and only enhancers with at least 10 CpG observations used for analysis. DMEs were defined within Seqmonk as enhancers that passed both a logistic regression test p≤0.01 and an EdgeR test p≤0.01, and had a minimum methylation difference of 30%.

CpG content was analysed against a background of mouse sequences using Compter (https://www.bioinformatics.babraham.ac.uk/projects/compter/). Motif analysis was performed on the central 500bp of DMEs using Homer (Heinz et al., 2010), against a background of all EpiCSeg enhancers. Cistrome DB Toolkit (Zheng et al., 2019) was used with default settings, meaning the top 1,000 peaks according to peak enrichment of each Cistrome sample were used for comparison, both for transcription factor and histone mark analysis. Genomic coordinates for the centre of mouse tissue-specific enhancers were sourced from Shen et al. (2012), ported from mm9 to mm10 using the UCSC Genome Browser LiftOver tool, and ±1.5kb windows around these were used for analysis. Annotations of Tet1 and Tet2 single knockout mESCs hyper/hypo-methylated DMRs were sourced from Hon et al. (2014).

*Published WGBS data used in this study:*

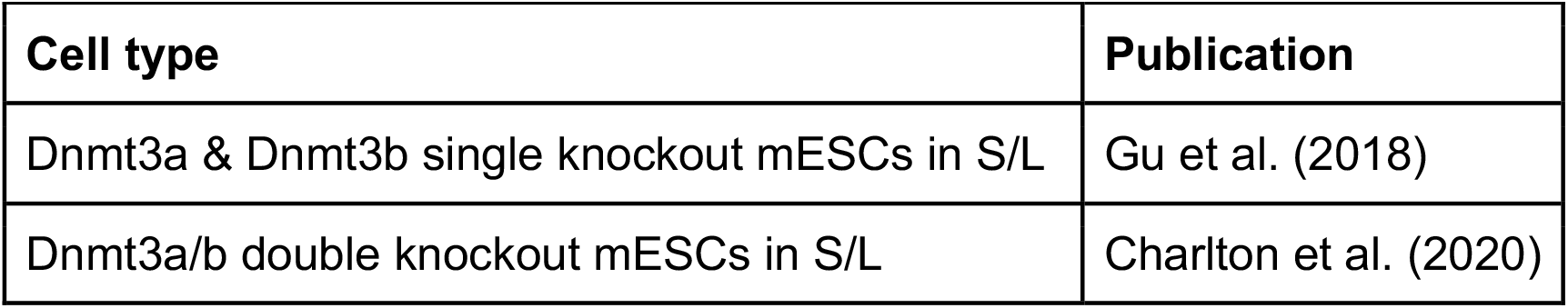

### In silico strand annealing

The methylation state of single CpG dyads was assessed by *in silico* strand annealing (iSA) as described in Xu and Corces (2019). Briefly, paired-end WGBS samples were processed using a standard Bismark workflow (as above - FastQ files trimmed with Trim Galore, aligned to the mouse GRCm38 genome using Bismark, and deduplicated with deduplicate_bismark). The deduplicated BAM files were used as input files for an interactive iSA shell script provided here: https://github.com/chxu02/iSA (sample command: “iSA.sh *deduplicated 0 0”; several commands had to be updated to work with a more recent version of bedtools; interactive prompts were removed). The total numbers of fully (un-)methylated or hemi-methylated dyads in CpG or CHG context were then summarised using a custom Python script.

### Hemimethylation patterning analysis

Starting from top/Watson and bottom/Crick strand iSA bam files, methylation information was extracted using Bismark Methylation Extractor. Using Seqmonk, hemimethylated CpG dyads were identified from opposing methylation calls on consecutive CG bases. Reads were included in the analysis if they contained at least three hemimethylated positions, and methylation calls for each of their CpGs were reported (custom Python script). Simulated data were obtained by first generating one million 200bp long genomic single-end top strand reads using a modified version of Sherman (https://github.com/FelixKrueger/Sherman). From these and their reverse complement counterparts, sequences were generated in which every C has a 50% chance of being converted to T. Downstream of this, simulated iSA reads were analysed in the same way as actual iSA reads.

### ChIP-seq and ATAC-seq analysis

Publicly available ChIP-seq and ATAC-seq data was trimmed to remove poor quality reads, adaptor and barcode sequences using Trim Galore. Trimmed data was mapped to the mouse GRCm38 genome assembly using Bowtie2 (Langmead and Salzberg, 2012), and mappings with MAPQ scores <20 were discarded. Mapped ChIP-seq and ATAC-seq data was imported into Seqmonk, with single-end reads extended by 200bp upon import. Transcription factor peaks were called within Seqmonk using a MACS peak caller (Zhang et al., 2008) with an uncorrected p-value of 1e-5 and average sonicated fragment size of 300bp. Peaks were called for replicate samples individually, using an appropriate input sample if available, then peak sets were combined across replicates. The TET2 peak set used was a combined set of peaks from TET2 ChIP-seq and TET2-FLAG ChIP-seq, excluding any peaks that overlapped those detected by TET2 ChIP-seq in Tet2 knockout mESCs, and FLAG ChIP-seq in a parental cell line in which TET2 was not FLAG-tagged.

Histone mark, TET2 and ATAC-seq signal over enhancers were quantitated within Seqmonk using a Read Count Quantitation normalising to enhancer length, thus generating log2 RPKM values (reads per kilobase, per million mapped reads of library). For histone mark scatter plots (Fig. 3C), enhancers with input ChIP-seq log2 RPKM values >2 were excluded from analysis.

*Published ChIP-seq / ATAC-seq data used in this study:*

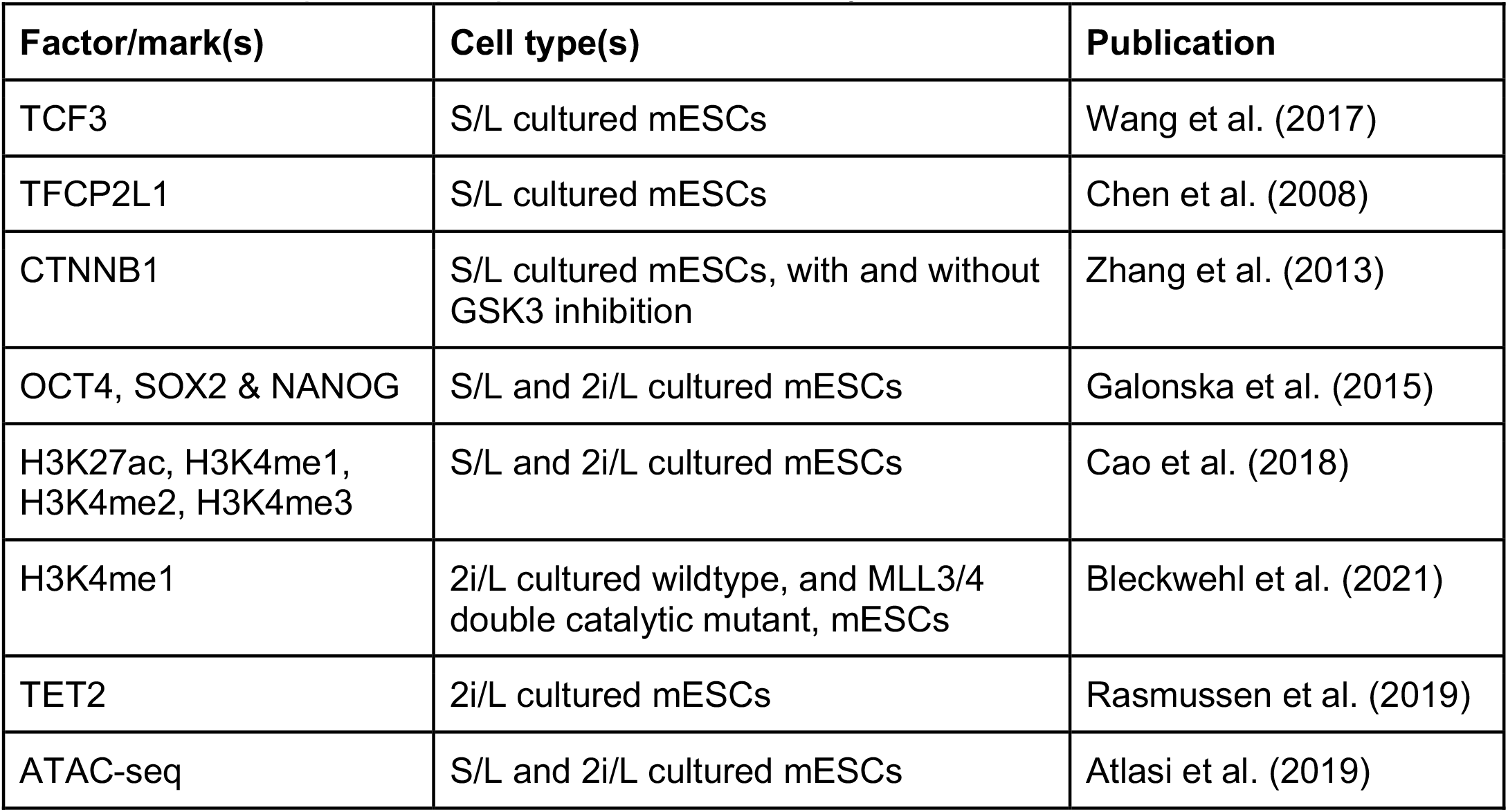

### Linking promoters with enhancers using CHi-C data

Published Capture Hi-C (CHi-C) interaction data from Atlasi et al. (2019) was used to link promoter with enhancer regions in the following way: Atlasi et al. report significant interactions between DNaseI sensitive regions for mESCs cultured in either S/L or 2i/L. Coordinates of all interacting regions were ported from mm9 to mm10 using the UCSC Genome Browser LiftOver tool irrespective of which culture conditions they originated in. To identify promoter-enhancer pairs, interacting regions were overlapped with promoter regions (-2kb to +500 bp of mRNA producing genes) and enhancer regions (EpiCSeq annotation from Peng et al. [2020]) using R packages GenomicRanges (Lawrence et al., 2013) and Splicejam (https://github.com/jmw86069/splicejam). Interactions were retained if they overlapped with at least one promoter and one enhancer either end.

#### RNA-seq analysis

Raw, single-end RNA-seq reads were trimmed using Trim Galore (v.0.6.5) with default settings. Trimmed reads were mapped to the mouse GRCm38 genome assembly using HISAT2 (Kim et al., 2019), and mappings with MAPQ scores <20 were discarded. Mapped RNA-seq data was quantitated over strand-specific mRNA probes using the Seqmonk RNA-seq Quantitation Pipeline to generate log2 RPM (reads per million mapped reads of library) expression values.

Genes were considered to be differentially expressed if they were significant (p<0.01) according to a DESeq2 Wald test.

## Author contributions

J.S. conceived the project, designed and performed experiments, analysed data and wrote the manuscript. C.K. analysed CpG hemimethylation patterning, created the EpiCSeg Shiny app used, analysed CpG methylation around hemimethylated CHG sites, created CHi-C linkage annotations, and provided assistance with and feedback on bioinformatic analyses, as well as on the manuscript. F.K. helped with initial feasibility assessments for WGBS, mapped and processed all sequencing data, and conducted *in silico* strand annealing analysis including patterning analysis. E.K.P. processed and ran all TMT mass spectrometry samples, and provided helpful input on experimental design and data interpretation. K.K. provided assistance with, and wrote code for, analysis of whole proteome data. C.S.D. supervised the proteomics experiment, providing helpful input on experimental design and data interpretation. W.R. supervised the study and provided expertise and feedback.

## Acknowledgements

We thank all members of the W.R. laboratory for helpful discussions, as well as members of Simon Cook’s laboratory for discussions on MEK signalling. We thank Stephen Clark, Stephen Bevan and Tim Lohoff for advice on WGBS protocols. We thank Nicole Forrester and Paula Kokko-Gonzales from the Babraham Institute NGS Facility, as well as Vicki Murray and Maike Paramor from the Cambridge Stem Cell Institute NGS Facility, for their help with generating and sequencing libraries. J.S. was supported by a Wellcome Trust PhD studentship (109140/Z/15/Z) and is currently supported by a Wellcome Trust – Investigator award (210754/Z/18/Z). Research in W.R.’s lab is supported by the BBSRC (BBS/E/B/000C0422) and Wellcome Trust – Investigator award (210754/Z/18/Z). W.R. is a consultant and shareholder of Cambridge Epigenetix.

---

**Figure S1:**
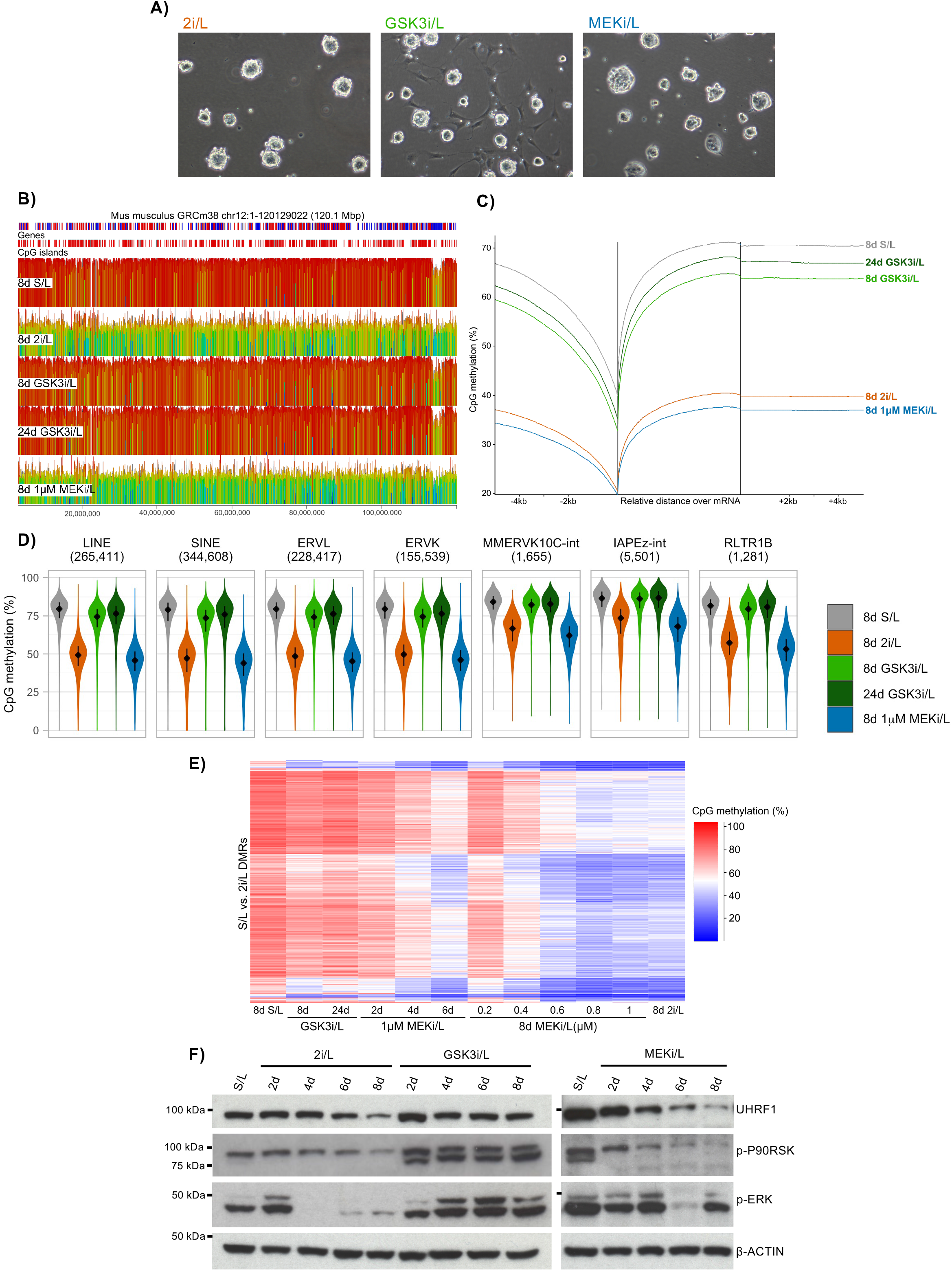
MEK inhibition causes dose-dependent impairment of maintenance methylation. **A)** Bright field microscopy images of E14Tg2a (XY) mouse embryonic stem cells grown in 2i/L, GSK3i/L or MEKi/L for 8 days. **B)**Percentage DNA methylation of 100 CpG windows over the entirety of chromosome 12, across culture conditions. Both vertical height and colour (from blue to red) of bars show percentage methylation. Gene annotations are coloured by strand (forward red; reverse blue). **C)**Percentage DNA methylation upstream of, over, and downstream of genes, across culture conditions. Percentage methylation was quantified over 100 CpG windows with at least 20 observations. **D)** Violin plots depicting percentage DNA methylation levels of different classes of repeat elements, across culture conditions. Percentage methylation was quantified over 100 CpG windows with at least 20 observations that overlap LINEs, SINEs, ERVL elements, ERVK elements, MMERVK10C-int elements (ERVK class), IAPEz-int elements (ERVK class), and RLTR1B elements (ERV1 class). The panels are labelled with the number of windows in each group. **E)** Heatmap depicting percentage DNA methylation of 2,500 randomly subsetted S/L vs. 2i/L DMRs across culture conditions. **F)** Western blot showing the abundance of UHRF1, phosphorylated p90 RSK, and phosphorylated ERK proteins across the 8d time course transition of S/L cultured cells into 2i/L, GSK3i/L or MEKi/L media. β-ACTIN quantity is shown as a loading control.

**Figure S2:**
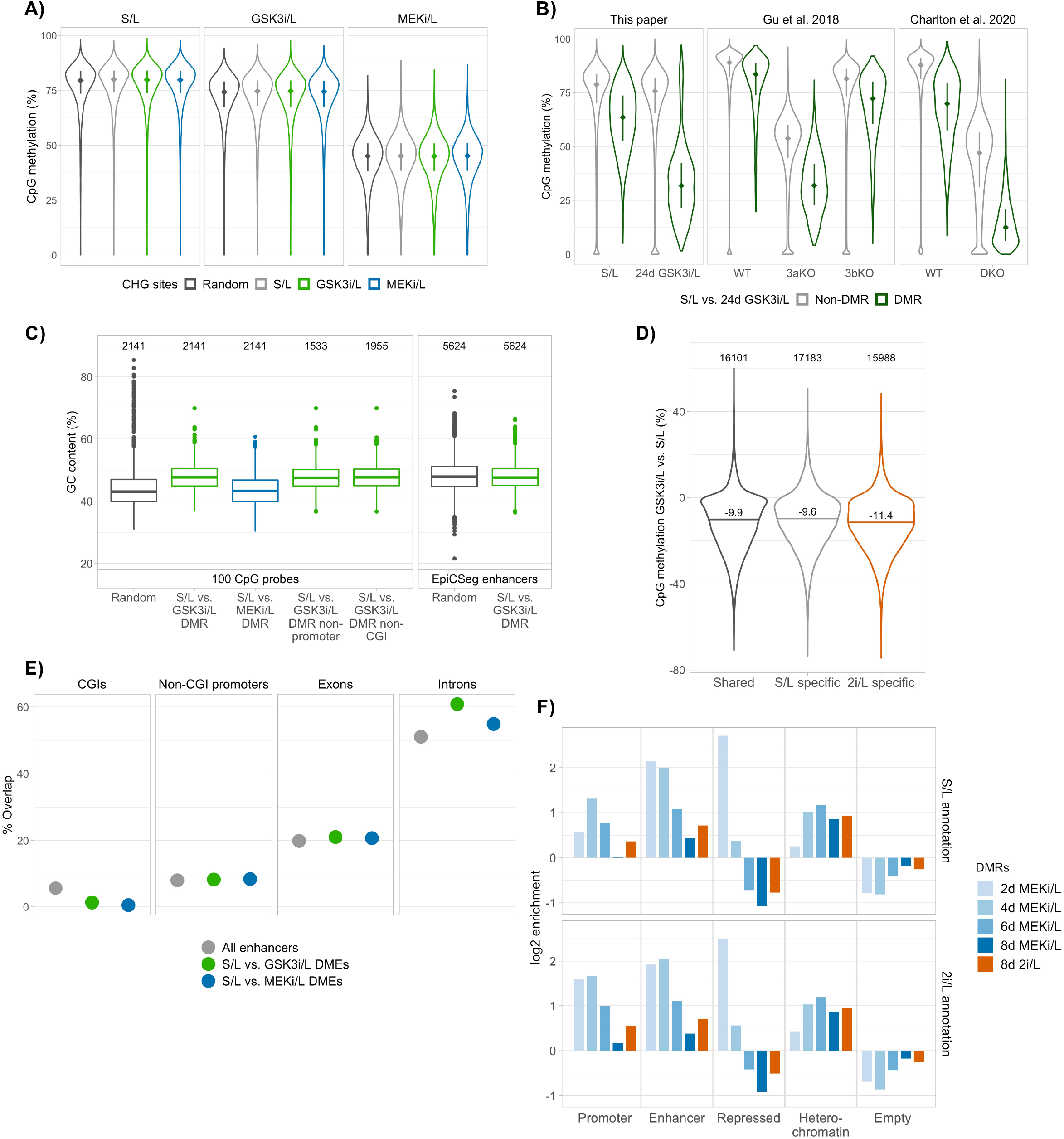
GSK3i/L-driven impairment of *de novo* methylation results in hypomethylation of enhancers and non-CGI promoters. **A)** Percentage CpG methylation of 100 CpG probes containing hemimethylated CHG sites in S/L (19,590 probes), GSK3i/L (32,785 probes) or MEKi/L (24,459 probes) versus those containing random CHG sites (25,611 probes), shown across S/L, GSK3i/L and MEKi/L cultured mESCs. **B)**Violin plots depicting percentage CpG methylation levels of S/L vs. 24d GSK3i/L non-DMRs (420,227) and DMRs (1,956), shown across S/L and 24d GSK3i/L samples, Dnmt3a and Dnmt3b single knockout mESCs, and Dnmt3a/3b double knockout mESCs (DKO). **C)**Boxplots illustrating GC content of random subsets of, as well as differentially methylated, 100 CpG probes and EpiCSeg enhancers, calculated using Compter. Also plotted for 100 CpG probes are the GSK3i/L DMRs with those overlapping -5000bp to +500bp promoter regions, or alternatively those overlapping CGIs, removed. The number of regions in each group is printed above the boxes. The random subsets, as well as the MEKi/L 100 CpG DMRs, were randomly downsampled to match the set size of the GSK3i/L DMRs in each case. **D)** Percentage CpG methylation change between S/L and GSK3i/L of EpiCSeg enhancers annotated in S/L that do not overlap those annotated in 2i/L, enhancers in 2i/L that do not overlap those in S/L, and enhancers that overlap across the two conditions. The number of enhancers in each group is printed above the violins. **E)**Dotplot depicting percentages of EpiCSeg enhancers that overlap CGIs, non-CGI promoters, exons and introns, shown for all enhancers, S/L vs. GSK3i/L DMEs and S/L vs. MEKi/L DMEs. **F)** log2 enrichments of DMRs between S/L and each MEKi/L time point for different classes of genomic element as defined by EpiCSeg (Epigenome Count-based Segmentation) based on S/L and 2i/L cultured mESCs.

**Figure S3:**
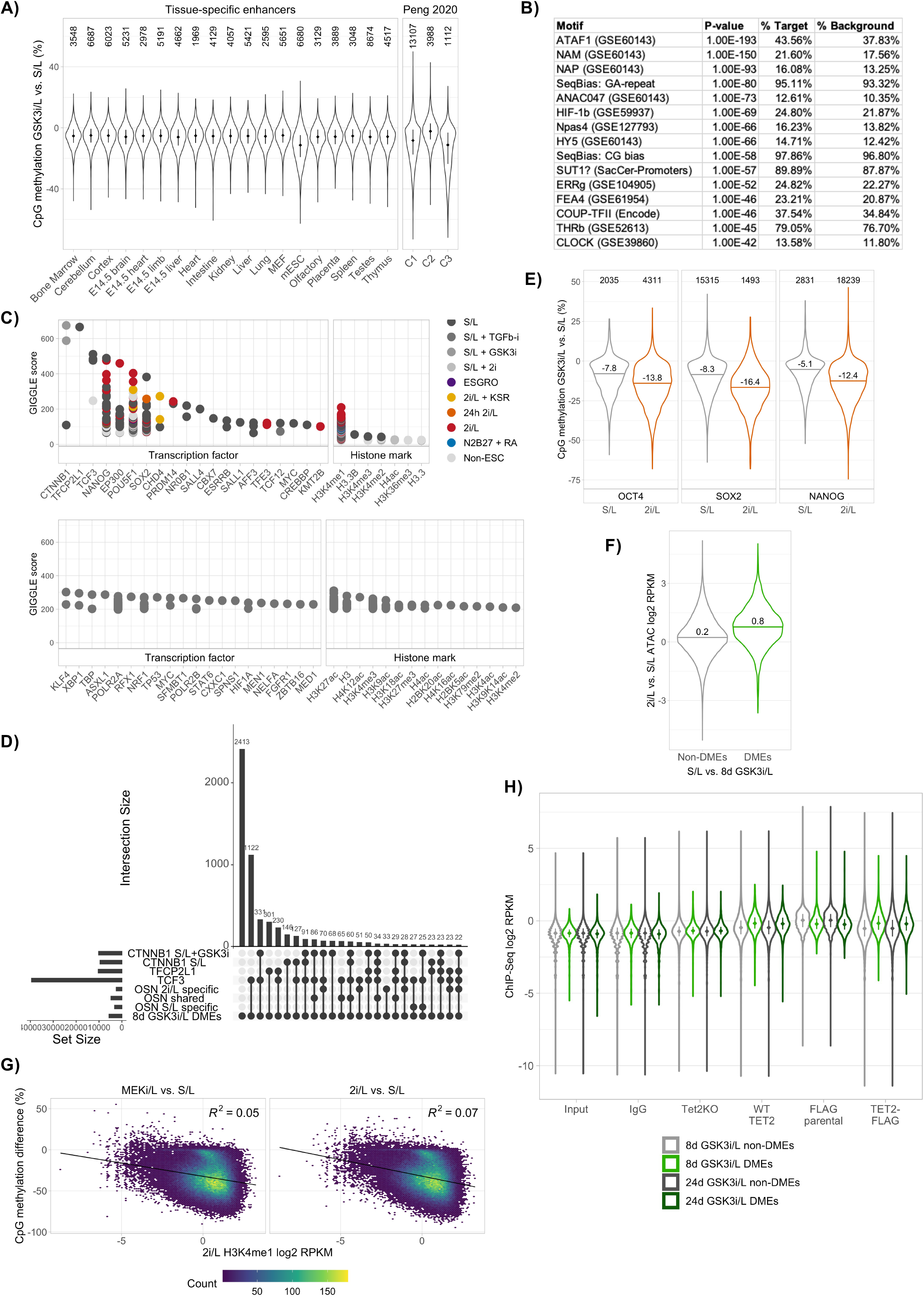
GSK3i/L hypomethylated enhancers are enriched for 2i/L-specific pluripotency factor binding, TET2 and H3K4me1. **A)** Violin plots depicting percentage CpG methylation change between S/L and 8d GSK3i/L at EpiCSeg enhancers overlapping different tissue-specific enhancers, or classes of enhancer as annotated by Peng et al. (2020). C1 enhancers were those with STARR-seq activity and active chromatin marks in S/L and/or 2i/L, C2 only STARR-seq activity, and C3 only active chromatin marks. The number of enhancers in each class is printed above the violins. **B)**The top results of Homer motif analysis for differentially methylated enhancers (DMEs) between S/L and 8d 1µM MEKi/L. **C)**Upper – the top results from Cistrome DB Toolkit, which quantifies overlap between ChIP-seq datasets in the database and genomic regions of interest, for S/L versus 8d GSK3i/L DMEs, coloured by culture condition of the Cistrome ChIP-seq datasets. Lower - top results from Cistrome DB Toolkit for S/L versus 8d GSK3i/L non-DMEs, not colour coded. **D)** Upset plot illustrating the degree of overlap between 8d GSK3i/L DMEs and selected transcription factor ChIP-seq peaks. The plot is 8d GSK3i/L DME centric, i.e. shows the top largest overlaps involving 8d GSK3i/L DMEs. Each overlap quantified is a distinct overlap, meaning the regions in that overlap do not also overlap with any other set of regions amongst those analysed. **E)** Violin plots depicting percentage CpG methylation change between S/L and 8d GSK3i/L at EpiCSeg enhancers overlapping OCT4, SOX2 or NANOG ChIP-seq peaks that were specific to either S/L or 2i/L cultured mESCs. The number of enhancers in each class is printed above the violins. **F)** Violin plots depicting ATAC-seq signal in 2i/L versus S/L at 8d GSK3i/L DMEs (5,624) or non- DMEs (178,511). **G)** Scatter plots coloured by density showing the relationship at EpiCSeg enhancers between percentage CpG methylation change in either 8d 2i/L or 1µM MEKi/L cultured mESCs compared to S/L, and H3K4me1 abundance in 2i/L. **H)** TET2 and TET2-FLAG binding in 2i/L at 8d and 24d GSK3i/L DMEs (5,624 and 4,650 respectively) versus non-DMEs (178,511 and 179,485 respectively). Also shown as controls – input signal, IgG binding, TET2 ChIP-seq in Tet2 knockout mESCs, and FLAG ChIP-seq in a parental cell line in which TET2 was not tagged with FLAG.

**Figure S4:**
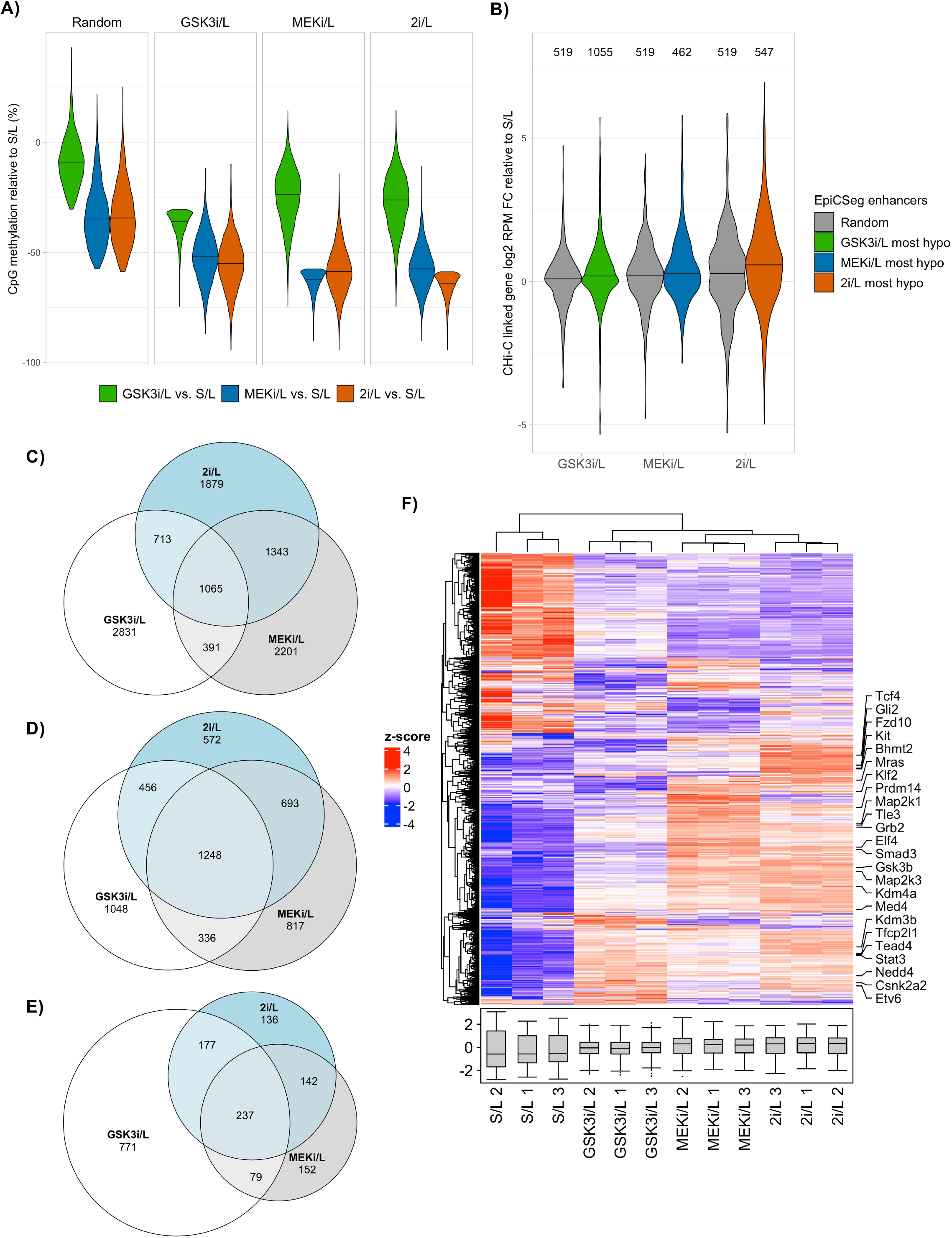
Enhancer hypomethylation coincides with upregulation of proximal genes. **A)** Violin plots depicting percentage CpG methylation change between S/L and GSK3i/L, MEKi/L or 2i/L of either random enhancers, or the 5,000 most hypomethylated enhancers in either GSK3i/L, MEKi/L or 2i/L, compared to S/L. These 5,000 enhancers were selected having pre- excluded enhancers overlapping -5000bp to +500bp promoter regions. Random enhancers are 5,000 randomly selected enhancers (not overlapping promoters) that are not amongst the 5,000 most hypomethylated in GSK3i/L, MEKi/L or 2i/L. **B)**Violin plots depicting gene expression (log2 RPM) fold change between S/L and GSK3i/L, MEKi/L or 2i/L. Expression is plotted of genes that are linked by Capture Hi-C (CHi-C) to either random enhancers, or to the 5,000 most hypomethylated enhancers (not overlapping promoters) in either GSK3i/L, MEKi/L or 2i/L, compared to S/L. The enhancer methylation comparisons correspond to the gene expression comparisons, i.e. linked gene expression in GSK3i/L compared to S/L is plotted for the 5,000 most hypomethylated enhancers in GSK3i/L compared to S/L. **C)**Euler diagram of the 5,000 most hypomethylated enhancers in either GSK3i/L, MEKi/L or 2i/L, compared to S/L. **D)** Euler diagram of the genes that are closest within 100kb to the 5,000 most hypomethylated enhancers in either GSK3i/L, MEKi/L or 2i/L, compared to S/L. **E)** Euler diagram of the genes that are linked by CHi-C to the 5,000 most hypomethylated enhancers in either GSK3i/L, MEKi/L or 2i/L, compared to S/L. **F)** Heatmap of protein expression for genes that are closest within 100kb or linked by CHi-C to the 5,000 most hypomethylated enhancers in 2i/L. Expression is shown across biological triplicates of 8d S/L, GSK3i/L, MEKi/L and 2i/L culture conditions. Selected genes are labelled.

## Bibliography

Atlasi, Y., Megchelenbrink, W., Peng, T., Habibi, E., Joshi, O., Wang, S.-Y., Wang, C., Logie, C., Poser, I., Marks, H., et al. (2019). Epigenetic modulation of a hardwired 3D chromatin landscape in two naive states of pluripotency. Nat. Cell Biol. 21, 568–578.

Berrens, R. V., Andrews, S., Spensberger, D., Santos, F., Dean, W., Gould, P., Sharif, J., Olova, N., Chandra, T., Koseki, H., et al. (2017). An endosiRNA-Based Repression Mechanism Counteracts Transposon Activation during Global DNA Demethylation in Embryonic Stem Cells. Cell Stem Cell 21, 694-703.e7.

Bleckwehl, T., Crispatzu, G., Schaaf, K., Respuela, P., Bartusel, M., Benson, L., Clark, S. J., Dorighi, K. M., Barral, A., Laugsch, M., et al. (2021). Enhancer-associated H3K4 methylation safeguards in vitro germline competence. Nat. Commun. 12, 5771.

Cao, K., Collings, C. K., Morgan, M. A., Marshall, S. A., Rendleman, E. J., Ozark, P. A., Smith, E. R. and Shilatifard, A. (2018). An Mll4/COMPASS-Lsd1 epigenetic axis governs enhancer function and pluripotency transition in embryonic stem cells. Sci. Adv. 4, eaap8747.

Chambers, I., Silva, J., Colby, D., Nichols, J., Nijmeijer, B., Robertson, M., Vrana, J., Jones, K., Grotewold, L. and Smith, A. (2007). Nanog safeguards pluripotency and mediates germline development. Nature 450, 1230–1234.

Charlton, J., Jung, E. J., Mattei, A. L., Bailly, N., Liao, J., Martin, E. J., Giesselmann, P., Brändl, B., Stamenova, E. K., Müller, F.-J., et al. (2020). TETs compete with DNMT3 activity in pluripotent cells at thousands of methylated somatic enhancers. Nat. Genet. 52, 819–827.

Chen, T., Ueda, Y., Dodge, J. E., Wang, Z. and Li, E. (2003). Establishment and maintenance of genomic methylation patterns in mouse embryonic stem cells by Dnmt3a and Dnmt3b. Mol. Cell. Biol. 23, 5594–5605.

Chen, X., Xu, H., Yuan, P., Fang, F., Huss, M., Vega, V. B., Wong, E., Orlov, Y. L., Zhang, W., Jiang, J., et al. (2008). Integration of external signaling pathways with the core transcriptional network in embryonic stem cells. Cell 133, 1106–1117.

Choi, J., Huebner, A. J., Clement, K., Walsh, R. M., Savol, A., Lin, K., Gu, H., Di Stefano, B., Brumbaugh, J., Kim, S.-Y., et al. (2017). Prolonged Mek1/2 suppression impairs the developmental potential of embryonic stem cells. Nature 548, 219–223.

Evans, M. J. and Kaufman, M. H. (1981). Establishment in culture of pluripotential cells from mouse embryos. Nature 292, 154–156.

Felle, M., Joppien, S., Németh, A., Diermeier, S., Thalhammer, V., Dobner, T., Kremmer, E., Kappler, R. and Längst, G. (2011). The USP7/Dnmt1 complex stimulates the DNA methylation activity of Dnmt1 and regulates the stability of UHRF1. Nucleic Acids Res. 39, 8355–8365.

Ficz, G., Hore, T. A., Santos, F., Lee, H. J., Dean, W., Arand, J., Krueger, F., Oxley, D., Paul, Y.-L., Walter, J., et al. (2013). FGF signaling inhibition in ESCs drives rapid genome-wide demethylation to the epigenetic ground state of pluripotency. Cell Stem Cell 13, 351–359.

Fidalgo, M., Huang, X., Guallar, D., Sanchez-Priego, C., Valdes, V. J., Saunders, A., Ding, J., Wu, W.-S., Clavel, C. and Wang, J. (2016). Zfp281 coordinates opposing functions of tet1 and tet2 in pluripotent states. Cell Stem Cell 19, 355–369.

Galonska, C., Ziller, M. J., Karnik, R. and Meissner, A. (2015). Ground State Conditions Induce Rapid Reorganization of Core Pluripotency Factor Binding before Global Epigenetic Reprogramming. Cell Stem Cell 17, 462–470.

Gu, T., Lin, X., Cullen, S. M., Luo, M., Jeong, M., Estecio, M., Shen, J., Hardikar, S., Sun, D., Su, J., et al. (2018). DNMT3A and TET1 cooperate to regulate promoter epigenetic landscapes in mouse embryonic stem cells. Genome Biol. 19, 88.

Habibi, E., Brinkman, A. B., Arand, J., Kroeze, L. I., Kerstens, H. H. D., Matarese, F., Lepikhov, K., Gut, M., Brun-Heath, I., Hubner, N. C., et al. (2013). Whole-genome bisulfite sequencing of two distinct interconvertible DNA methylomes of mouse embryonic stem cells. Cell Stem Cell 13, 360–369.

Hackett, J. A., Dietmann, S., Murakami, K., Down, T. A., Leitch, H. G. and Surani, M. A. (2013). Synergistic mechanisms of DNA demethylation during transition to ground-state pluripotency. Stem Cell Reports 1, 518–531.

Hayashi, K., de Sousa Lopes, S.M.C., Tang, F., Lao, K. and Surani, M. A. (2008). Dynamic equilibrium and heterogeneity of mouse pluripotent stem cells with distinct functional and epigenetic states. Cell Stem Cell 3, 391–401.

Heinz, S., Benner, C., Spann, N., Bertolino, E., Lin, Y. C., Laslo, P., Cheng, J. X., Murre, C., Singh, H. and Glass, C. K. (2010). Simple combinations of lineage-determining transcription factors prime cis-regulatory elements required for macrophage and B cell identities. Mol. Cell 38, 576–589.

Hon, G. C., Song, C.-X., Du, T., Jin, F., Selvaraj, S., Lee, A. Y., Yen, C.-A., Ye, Z., Mao, S.-Q., Wang, B.-A., et al. (2014). 5mC oxidation by Tet2 modulates enhancer activity and timing of transcriptome reprogramming during differentiation. Mol. Cell 56, 286–297.

Jeong, M., Sun, D., Luo, M., Huang, Y., Challen, G. A., Rodriguez, B., Zhang, X., Chavez, L., Wang, H., Hannah, R., et al. (2014). Large conserved domains of low DNA methylation maintained by Dnmt3a. Nat. Genet. 46, 17–23.

Kim, D., Paggi, J. M., Park, C., Bennett, C. and Salzberg, S. L. (2019). Graph-based genome alignment and genotyping with HISAT2 and HISAT-genotype. Nat. Biotechnol. 37, 907–915.

Kobayashi, H., Sakurai, T., Miura, F., Imai, M., Mochiduki, K., Yanagisawa, E., Sakashita, A., Wakai, T., Suzuki, Y., Ito, T., et al. (2013). High-resolution DNA methylome analysis of primordial germ cells identifies gender-specific reprogramming in mice. Genome Res. 23, 616–627.

Krueger, F. and Andrews, S. R. (2011). Bismark: a flexible aligner and methylation caller for Bisulfite-Seq applications. Bioinformatics 27, 1571–1572.

Kunath, T., Saba-El-Leil, M. K., Almousailleakh, M., Wray, J., Meloche, S. and Smith, A. (2007). FGF stimulation of the Erk1/2 signalling cascade triggers transition of pluripotent embryonic stem cells from self-renewal to lineage commitment. Development 134, 2895–2902.

Langmead, B. and Salzberg, S. L. (2012). Fast gapped-read alignment with Bowtie 2. Nat. Methods 9, 357–359.

Lawrence, M., Huber, W., Pagès, H., Aboyoun, P., Carlson, M., Gentleman, R., Morgan, M. T. and Carey, V. J. (2013). Software for computing and annotating genomic ranges. PLoS Comput. Biol. 9, e1003118.

Leitch, H. G., McEwen, K. R., Turp, A., Encheva, V., Carroll, T., Grabole, N., Mansfield, W., Nashun, B., Knezovich, J. G., Smith, A., et al. (2013). Naive pluripotency is associated with global DNA hypomethylation. Nat. Struct. Mol. Biol. 20, 311–316.

Marks, H., Kalkan, T., Menafra, R., Denissov, S., Jones, K., Hofemeister, H., Nichols, J., Kranz, A., Stewart, A. F., Smith, A., et al. (2012). The transcriptional and epigenomic foundations of ground state pluripotency. Cell 149, 590–604.

Martin, G. R. (1981). Isolation of a pluripotent cell line from early mouse embryos cultured in medium conditioned by teratocarcinoma stem cells. Proc Natl Acad Sci USA 78, 7634–7638.

Nichols, J., Silva, J., Roode, M. and Smith, A. (2009). Suppression of Erk signalling promotes ground state pluripotency in the mouse embryo. Development 136, 3215–3222.

Ooi, S. K. T., Qiu, C., Bernstein, E., Li, K., Jia, D., Yang, Z., Erdjument-Bromage, H., Tempst, P., Lin, S.-P., Allis, C. D., et al. (2007). DNMT3L connects unmethylated lysine 4 of histone H3 to de novo methylation of DNA. Nature 448, 714–717.

Papachristou, E. K., Kishore, K., Holding, A. N., Harvey, K., Roumeliotis, T. I., Chilamakuri, C. S. R., Omarjee, S., Chia, K. M., Swarbrick, A., Lim, E., et al. (2018). A quantitative mass spectrometry-based approach to monitor the dynamics of endogenous chromatin-associated protein complexes. Nat. Commun. 9, 2311.

Peng, T., Zhai, Y., Atlasi, Y., Ter Huurne, M., Marks, H., Stunnenberg, H. G. and Megchelenbrink, W. (2020). STARR-seq identifies active, chromatin-masked, and dormant enhancers in pluripotent mouse embryonic stem cells. Genome Biol. 21, 243.

Perez-Riverol, Y., Csordas, A., Bai, J., Bernal-Llinares, M., Hewapathirana, S., Kundu, D. J., Inuganti, A., Griss, J., Mayer, G., Eisenacher, M., et al. (2019). The PRIDE database and related tools and resources in 2019: improving support for quantification data. Nucleic Acids Res. 47, D442–D450.

Petell, C. J., Alabdi, L., He, M., San Miguel, P., Rose, R. and Gowher, H. (2016). An epigenetic switch regulates de novo DNA methylation at a subset of pluripotency gene enhancers during embryonic stem cell differentiation. Nucleic Acids Res. 44, 7605–7617.

Rasmussen, K. D., Berest, I., Keβler, S., Nishimura, K., Simón-Carrasco, L., Vassiliou, G. S., Pedersen, M. T., Christensen, J., Zaugg, J. B. and Helin, K. (2019). TET2 binding to enhancers facilitates transcription factor recruitment in hematopoietic cells. Genome Res. 29, 564–575.

Rulands, S., Lee, H. J., Clark, S. J., Angermueller, C., Smallwood, S. A., Krueger, F., Mohammed, H., Dean, W., Nichols, J., Rugg-Gunn, P., et al. (2018). Genome-Scale Oscillations in DNA Methylation during Exit from Pluripotency. Cell Syst. 7, 63-76.e12.

Seisenberger, S., Andrews, S., Krueger, F., Arand, J., Walter, J., Santos, F., Popp, C., Thienpont, B., Dean, W. and Reik, W. (2012). The dynamics of genome-wide DNA methylation reprogramming in mouse primordial germ cells. Mol. Cell 48, 849–862.

Sharifi-Zarchi, A., Gerovska, D., Adachi, K., Totonchi, M., Pezeshk, H., Taft, R. J., Schöler, H. R., Chitsaz, H., Sadeghi, M., Baharvand, H., et al. (2017). DNA methylation regulates discrimination of enhancers from promoters through a H3K4me1-H3K4me3 seesaw mechanism. BMC Genomics 18, 964.

Shen, Y., Yue, F., McCleary, D. F., Ye, Z., Edsall, L., Kuan, S., Wagner, U., Dixon, J., Lee, L., Lobanenkov, V. V., et al. (2012). A map of the cis-regulatory sequences in the mouse genome. Nature 488, 116–120.

Sim, Y.-J., Kim, M.-S., Nayfeh, A., Yun, Y.-J., Kim, S.-J., Park, K.-T., Kim, C.-H. and Kim, K.-S. (2017). 2i Maintains a Naive Ground State in ESCs through Two Distinct Epigenetic Mechanisms. Stem Cell Reports 8, 1312–1328.

Smith, A. G., Heath, J. K., Donaldson, D. D., Wong, G. G., Moreau, J., Stahl, M. and Rogers, D. (1988). Inhibition of pluripotential embryonic stem cell differentiation by purified polypeptides. Nature 336, 688–690.

Takashima, Y., Guo, G., Loos, R., Nichols, J., Ficz, G., Krueger, F., Oxley, D., Santos, F., Clarke, J., Mansfield, W., et al. (2014). Resetting transcription factor control circuitry toward ground-state pluripotency in human. Cell 158, 1254–1269.

Toyooka, Y., Shimosato, D., Murakami, K., Takahashi, K. and Niwa, H. (2008). Identification and characterization of subpopulations in undifferentiated ES cell culture. Development 135, 909–918.

von Meyenn, F., Iurlaro, M., Habibi, E., Liu, N. Q., Salehzadeh-Yazdi, A., Santos, F., Petrini, E., Milagre, I., Yu, M., Xie, Z., et al. (2016). Impairment of DNA methylation maintenance is the main cause of global demethylation in naive embryonic stem cells. Mol. Cell 62, 848–861.

Wang, Q., Zou, Y., Nowotschin, S., Kim, S. Y., Li, Q. V., Soh, C.-L., Su, J., Zhang, C., Shu, W., Xi, Q., et al. (2017). The p53 family coordinates wnt and nodal inputs in mesendodermal differentiation of embryonic stem cells. Cell Stem Cell 20, 70–86.

Wray, J., Kalkan, T. and Smith, A. G. (2010). The ground state of pluripotency. Biochem. Soc. Trans. 38, 1027–1032.

Wray, J., Kalkan, T., Gomez-Lopez, S., Eckardt, D., Cook, A., Kemler, R. and Smith, A. (2011). Inhibition of glycogen synthase kinase-3 alleviates Tcf3 repression of the pluripotency network and increases embryonic stem cell resistance to differentiation. Nat. Cell Biol. 13, 838–845.

Xu, C. and Corces, V. G. (2019). Resolution of the DNA methylation state of single CpG dyads using in silico strand annealing and WGBS data. Nat. Protoc. 14, 202–216.

Yamaji, M., Ueda, J., Hayashi, K., Ohta, H., Yabuta, Y., Kurimoto, K., Nakato, R., Yamada, Y., Shirahige, K. and Saitou, M. (2013). PRDM14 ensures naive pluripotency through dual regulation of signaling and epigenetic pathways in mouse embryonic stem cells. Cell Stem Cell 12, 368–382.

Ying, Q.-L., Wray, J., Nichols, J., Batlle-Morera, L., Doble, B., Woodgett, J., Cohen, P. and Smith, A. (2008). The ground state of embryonic stem cell self-renewal. Nature 453, 519–523.

Zhang, Y., Liu, T., Meyer, C. A., Eeckhoute, J., Johnson, D. S., Bernstein, B. E., Nusbaum, C., Myers, R. M., Brown, M., Li, W., et al. (2008). Model-based analysis of ChIP-Seq (MACS). Genome Biol. 9, R137.

Zhang, Y., Jurkowska, R., Soeroes, S., Rajavelu, A., Dhayalan, A., Bock, I., Rathert, P., Brandt, O., Reinhardt, R., Fischle, W., et al. (2010). Chromatin methylation activity of Dnmt3a and Dnmt3a/3L is guided by interaction of the ADD domain with the histone H3 tail. Nucleic Acids Res. 38, 4246–4253.

Zhang, X., Peterson, K. A., Liu, X. S., McMahon, A. P. and Ohba, S. (2013). Gene regulatory networks mediating canonical Wnt signal-directed control of pluripotency and differentiation in embryo stem cells. Stem Cells 31, 2667–2679.

Zheng, R., Wan, C., Mei, S., Qin, Q., Wu, Q., Sun, H., Chen, C.-H., Brown, M., Zhang, X., Meyer, C. A., et al. (2019). Cistrome Data Browser: expanded datasets and new tools for gene regulatory analysis. Nucleic Acids Res. 47, D729–D735.

